# Human primitive mesenchymal stem cell-derived retinal progenitor cells promoted neuroprotection and neurogenesis in rd12 mice

**DOI:** 10.1101/2021.09.20.460984

**Authors:** Christina Brown, Patrina Agosta, Christina McKee, Keegan Walker, Matteo Mazzella, David Svinarich, G. Rasul Chaudhry

## Abstract

Retinal degenerative diseases (RDD) such as retinitis pigmentosa (RP) have no treatment. Stem cell-based therapies could provide promising opportunities to repair the damaged retina and restore vision. We investigated a novel approach in which human retinal progenitor cells (RPCs) derived from primitive mesenchymal stem cells (pMSCs) were examined to treat retinal degeneration in an rd12 mouse model of RP. Intravitreally transplanted cells improved retinal function and significantly increased retinal thickness. Transplanted cells homed, survived, and integrated to various retinal layers. They also induced anti-inflammatory and neuroprotective responses and upregulated neurogenesis genes. We found that RPCs were more efficacious than pMSCs in improving the retinal structure and function. RNA analyses suggest that RPCs promote neuroprotection and neuronal differentiation by activating JAK/STAT and MAPK, and inhibiting BMP signaling pathways. These promising results provide the basis for clinical studies to treat RDD using RPCs derived from pMSCs.

## Introduction

Retinal degenerative diseases (RDD), including age-related macular degeneration (AMD) and retinitis pigmentosa (RP) cause progressive and incurable vision loss^[1, 2]^. AMD leads to the loss of sharp and central vision due to damage to the macula that affects the central area of the retina and choroid^[3]^. Whereas RP is characterized by progressive degeneration of the photoreceptors, leading to night blindness followed by progressive loss of the visual field in daylight and ultimately blindness^[4]^. RP affects 1 in every 4000 people in the United States and 1 in every 5000 people worldwide^[2]^. In RP, mutations in the gene, *RPE65*, results in the disruption of the RPE65 protein production in the retinal pigment epithelium (RPE)^[5]^, leading to worsened vision, night blindness, and decreased peripheral vision^[6]^. A mouse model, rd12, represents RP and Leber congenital amaurosis in humans, which causes blindness or severely impaired vision in adults and children, respectively^[7]^. The mutation responsible for vision loss in the rd12 model is a nonsense mutation within the *Rpe65* gene^[8]^. Using this animal model, we have previously demonstrated the therapeutic potential of embryonic stem cells (ESCs) derived neuroprogenitors^[9]^. Transplantation of pluripotent stem cells, ESC- and induced pluripotent stem cell (iPSC-) derived retinal progenitors have also been reported to preserve visual function upon injection into Royal College of Surgeons (RCS) rats as well as rd1 and rd12 mice, respectively^[10–12]^. Because pluripotent cells can cause teratomas and pose moral and ethical concerns, growing interest has been shown to explore other options such as mesenchymal stem cells (MSCs) for cell therapy.

MSCs exert their therapeutic effect in part by secreting trophic factors that protect cells or promote their survival and transplantation of mouse bone marrow (BM)-MSCs delayed retinal degeneration in an animal model of RDD^[13]^. Mouse BM-MSCs transplanted into rhodopsin knock-out mice integrated into the RPE and the neuroretina, which lead to prolonged photoreceptor survival^[14]^. Human (h) BM-MSCs injected into the subretinal space of a retinal dystrophy rat model showed significant and extensive photoreceptor rescue in transplanted eyes^[15]^. On the other hand, tail vein injection of rat MSCs preserved visual function in a rat model of RP^[16]^. Interestingly, transplantation of the combination of hBM-MSCs and fetal retinal progenitor cells (RPCs) into the subretinal space of a rat model of RP maintained retinal function based on the electroretinogram (ERG) results much better than when they were individually transplanted^[17]^. However, these findings are difficult to translate in humans, mainly because of the invasive procedures required to isolate these cells. In addition, adult MSCs have limited proliferation and differentiation potential^[18]^. Furthermore, adult MSCs undergo genetic changes depending on donor age and environmental stress exposure^[19]^. Recently, human umbilical cord (hUC) MSCs have been investigated for treating retinal degeneration. Human Wharton’s jelly (hWJ) MSCs provided neuroprotection for retinal ganglion cells (RGCs) and promoted axonal regeneration along the optic nerve in adult rats^[20]^. hWJ-MSCs also delayed loss of RGC in axotomy-induced rats^[21]^, outer nuclear layer (ONL) in RCS rats^[22]^, and improved the thickness of the retina and upregulated pro-survival genes^[23]^. Human umbilical cord blood MSCs were suggested to improve the retinal morphological structure^[24]^, exert a neuroprotective effect and rescue a significant percentage of axotomized RGCs^[25]^.

The potential therapeutic effects of hBM-MSCs or adipose MSCs on retinal defects have been investigated in 30 clinical trials^[26–29]^. In some MSC transplantation studies, vision loss was halted, macular thickness increased, and/or visual acuity was improved^[30–32]^. Human RPC transplantation in RP patients led to moderate vision recovery and maintenance of the ONL thickness^[33]^ or improved visual acuity^[34]^. It is important to note that most clinical trials are limited to phase I/II studies^[26, 35]^, due to the lack of cells required for extensive clinical studies.

The current study investigated human primitive (p) MSCs isolated from perinatal tissue, which exhibited superior characteristics compared to adult MSCs and differentiated into RPCs. When pMSC-derived RPCs were transplanted in the rd12 mice, they improved retinal structure and function. They also survived, dispersed, and integrated into various neural layers, while pMSCs homed to the RPE layer of the retina. These findings provide evidence that pMSC-derived RPCs are a promising source for cell therapy to treat RDD.

## Results

### Differentiation and characterization of pMSCs into RPCs

Using a differentiation medium, we used previously isolated and characterized pMSCs pMSCs^[36]^ to differentiate them into RPCs. The differentiated cells exhibited neural outgrowth and downregulated the mesenchymal stem cell markers, CD29, CD44, CD73, CD90, and CD105, but upregulated the protein RCVRN, neural genes *TUJ1, NESTIN,* and *PAX6,* and retinal genes, *RCVRN, CRX, RHO, SIX3*, *OTX2, SIX6, GNL3, FABP7, NF200, SSEA4, ABCG2, DACH1, PTK7, PCNA, β-CATENIN, NOTCH1,* and *RPS27A* (Fig. 1A-D). The expression of neural and retinal markers was further validated by immunofluorescence staining by antibodies against TUJ1, NESTIN, PAX6, RCVRN, CRX, and RHO (Fig.1E-G). Based on these results, the differentiated derivatives of pMSCs were considered to be RPCs. We estimated the rate of differentiation of pMSCs into RPCs was 83%, 91%, 89%, 90%, 94%, and 81% based on the expression of TUJ1, NESTIN, PAX6, RCVRN, CRX, and RHO, respectively. RPCs also expressed high levels of neurotrophic factors, *BDNF, GDNF, IGF, CNTF, PDGF, EGF,* and *FGF* compared to pMSCs (Fig. 1H), suggesting that they could potentially promote neuroprotection.

**Figure 1:**
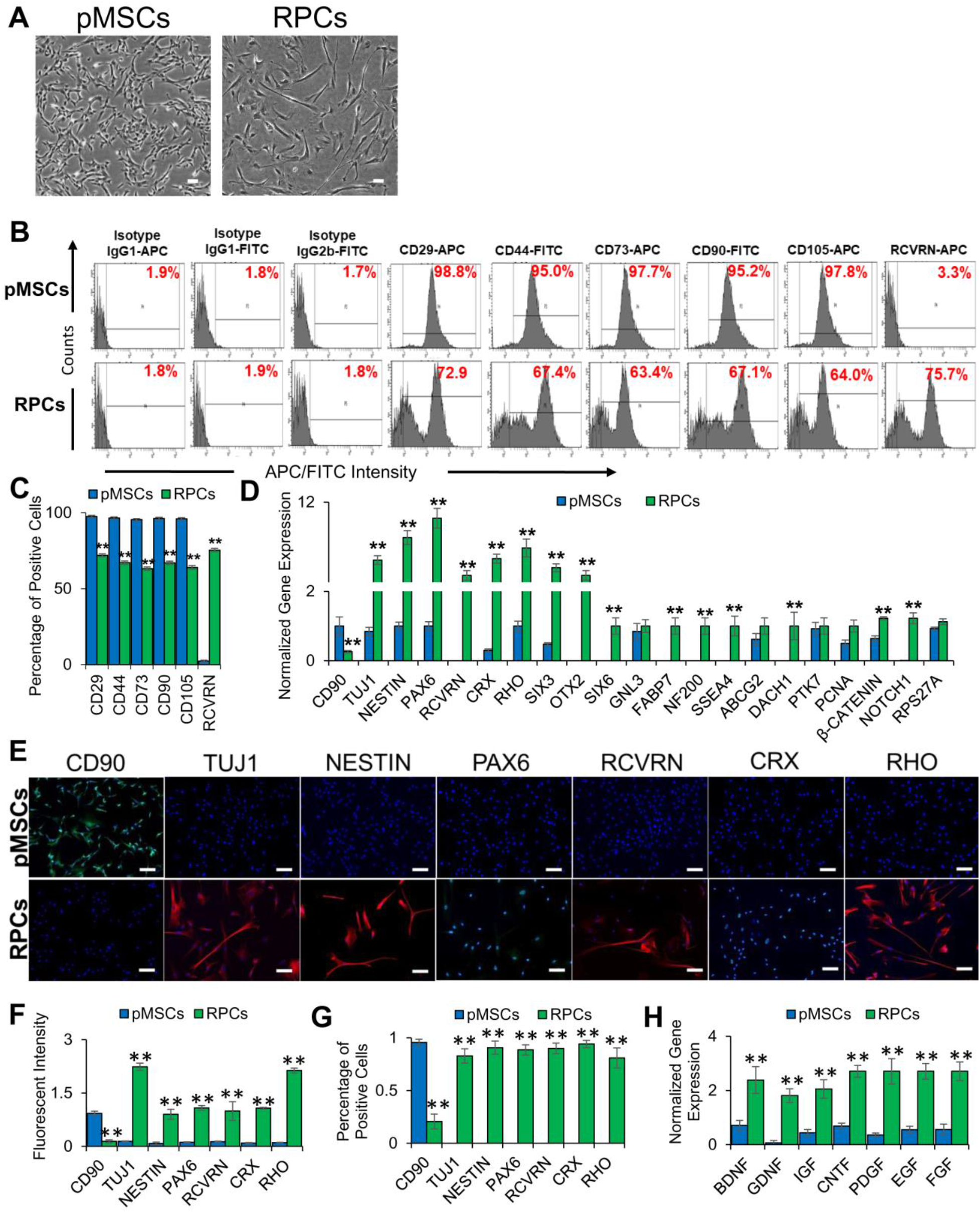
Characterization of RPCs derived from pMSCs. (A) Phase-contrast images of pMSCs and RPCs. 100 µm scale bar. (Magnification: 4x). (B and C) Histograms and graphical representation of the expression of MSC and retinal markers by pMSCs and RPCs as determined by flow cytometry (**p ≤ 0.01). (D) Expression of MSC (*CD90*), neural (*TUJ1, NESTIN,* and *PAX6*), retinal (*RCVRN, CRX, RHO, SIX3, OTX2, SIX6, GNL3, FABP7, NF200, SSEA4, ABCG2, DACH1, PTK7, PCNA, β-CATENIN, NOTCH1,* and *RPS27A*) genes in pMSCs and RPCs as determined by qRT-PCR. Gene expression was normalized to *GAPDH* and *β-ACTIN*. Error bars represent the SEM (**p ≤ 0.01). (E) Expression of MSC (CD90), neural (TUJ1, NESTIN, and PAX6), and retinal (RCVRN, CRX, and RHO) proteins in pMSCs and RPCs as visualized by immunocytostaining. Shown are representative merged images of DAPI (blue) and human antibodies (green and red). 100 µm scale bar (Magnification: 10x). (F) Measurement of fluorescent intensity of proteins immunocytostained using antibodies in pMSCs and RPCs (**p ≤ 0.01). (G) Percentage of positive immunocytostained cells (**p ≤ 0.01). (H) Expression of neurotropic genes, *BDNF, GDNF, IGF, CNTF, PDGF, EGF,* and *FGF* in pMSCs and RPCs. Gene expression was normalized to *GAPDH* and *β-ACTIN*, and error bars represent the SEM (**p ≤ 0.01). Significant changes in the gene and protein expression were observed upon differentiation of pMSCs into RPCs. All experiments were carried out in triplicates.

### Transcriptomic analysis of pMSCs and RPCs

To further characterize the RPCs, RNA-seq analysis was performed. The results depicted in Fig. 2A show a distinct difference in gene expression patterns between pMSCs and RPCs. This analysis revealed 2951 differentially expressed genes (DEGs) at a false discovery rate (FDR) < 0.05. Of these DEGs, 1,618 genes were upregulated in pMSCs, and 1,333 genes were upregulated in RPCs. Among the top 100 DEGs (FDR < 8.3 x 10^-147^), 68 and 32 genes were upregulated in pMSCs and RPCs, respectively (Fig. 2B-C). Notably, 8 upregulated DEGs in pMSCs were specific to RPE while 18 upregulated DEGs in RPCs were specific to the neuronal layers of the retina (FDR < 0.05) (Fig. 2D). The expression of selected DEGs was validated by quantitative reverse transcriptase-polymerase chain reaction (qRT-PCR). The selected DEGs were associated with retinal cells including, photoreceptors *(SULF2, GNAT2 and NRL),* amacrine *(TFAP2A and ATP1B1),* horizontal *(NDRG1),* Müller glia *(HES1, and SPON1),* and ganglion *(EBF1)* cells (Fig. 2E).

**Figure 2:**
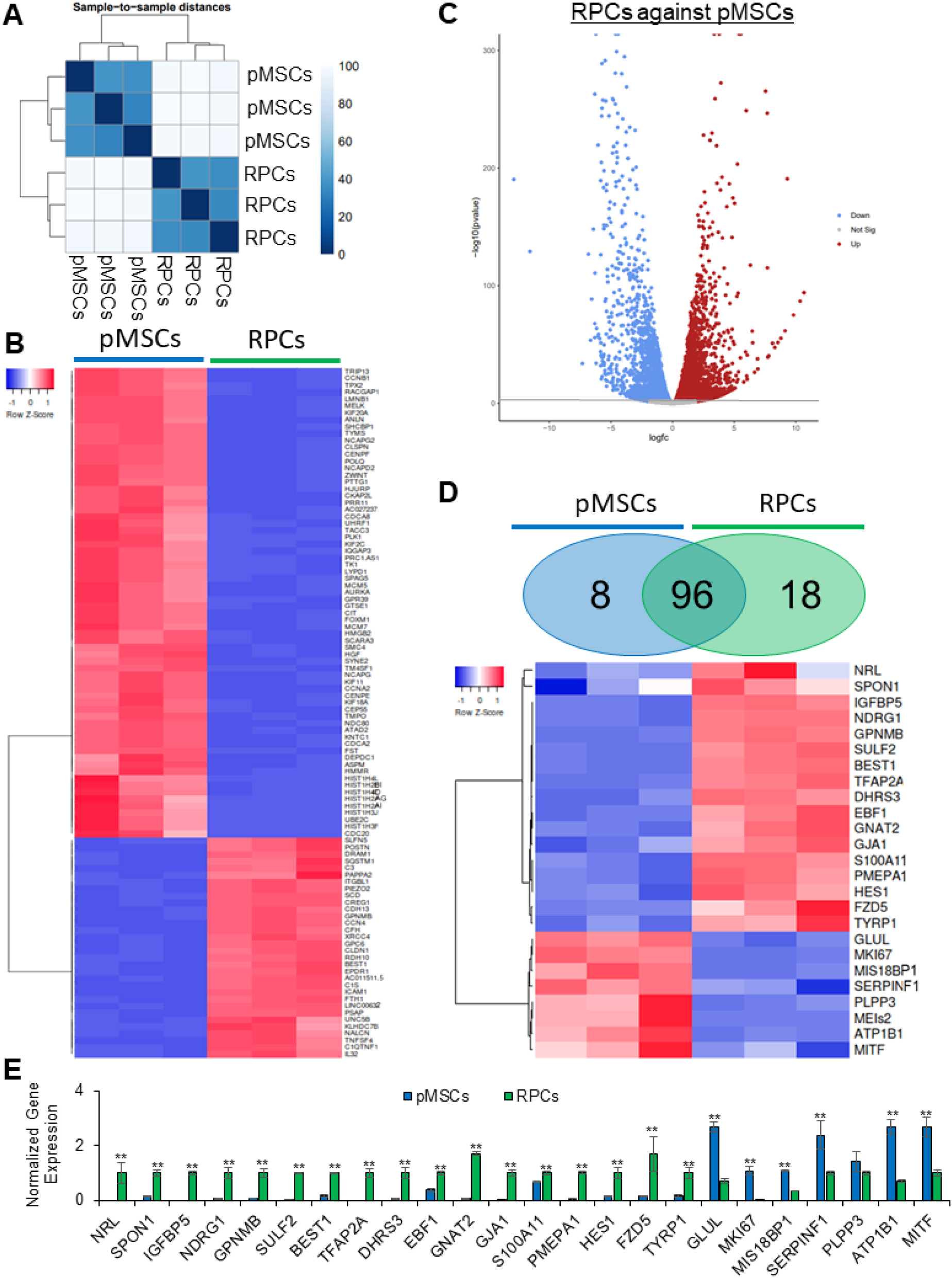
Transcriptome analysis of pMSCs and RPCs. (A) Hierarchical clustering analysis was performed using DESeq2, and the data was plotted as a sample-to-sample heatmap. The colored indicator represents sample distances (blue signifies a high correlation). (B) Heatmap showing raw z-scores of RNA-seq log2 transformed values of the top 100 DEGs and (C) volcano plot to visualize DEGs. (D) Heatmap showing raw z-scores of RNA-seq log2 transformed values of expression of retinal markers in pMSCs and RPCs. Venn diagrams depict the total number of genes upregulated in pMSCs (blue) and RPCs (green). In the heatmap, upregulated and downregulated genes are shown in red and blue, respectively. (E) Validation of selected retinal DEGs in pMSCs and RPCs by qRT-PCR. Gene expression was normalized to *GAPDH* and *β-ACTIN*, and error bars represent the SEM (**p ≤ 0.01). All experiments were performed in triplicate.

We next performed functional analysis of the DEGs using gene ontology (GO) analysis. Comparative enrichment analysis of pMSCs and RPCs performed using Enrichr identified DEGs associated with biological process, molecular function, and cellular component ontologies. In the category of biological processes, pMSCs were highly enriched in DNA ligation, chromosome organization, and positive regulation of cell cycle process (Fig. 3A). On the other hand, RPCs were significantly enriched in regulation of axon guidance, positive regulation of stem cell differentiation, dendritic spine maintenance, astrocyte activation, negative regulation of FGF receptor signaling pathway, and regulation of regulation mesenchymal stem cell differentiation, neuron projection maintenance, and extracellular matrix organization. Next, we investigated the category of molecular function (Fig. 3B). pMSCs were enriched in DNA replication origin binding, ATP-dependent DNA helicase activity, and 3’-5’ DNA helicase activity. Whereas RPCs presented a significant enrichment of integrin binding, sugar:proton symporter activity, low-density lipoprotein receptor activity, insulin-like growth factor I binding, serine-type carboxypeptidase activity, protein-lysine 6-oxidase activity, sulfuric ester hydrolase activity, and insulin-like growth factor II binding. We then investigated cellular components enriched in the DEGs. GO analysis also showed that chromatin, spindle microtubule, and condensed nuclear chromosome were enriched in pMSCs. While RPCs displayed significant enrichment of primary lysosome, tertiary granule lumen, dystrophin-associated glycoprotein complex, lytic vacuole, endoplasmic reticulum lumen, lysosome, vacuolar lumen, and secondary lysosome (Fig. 3C).

**Figure 3:**
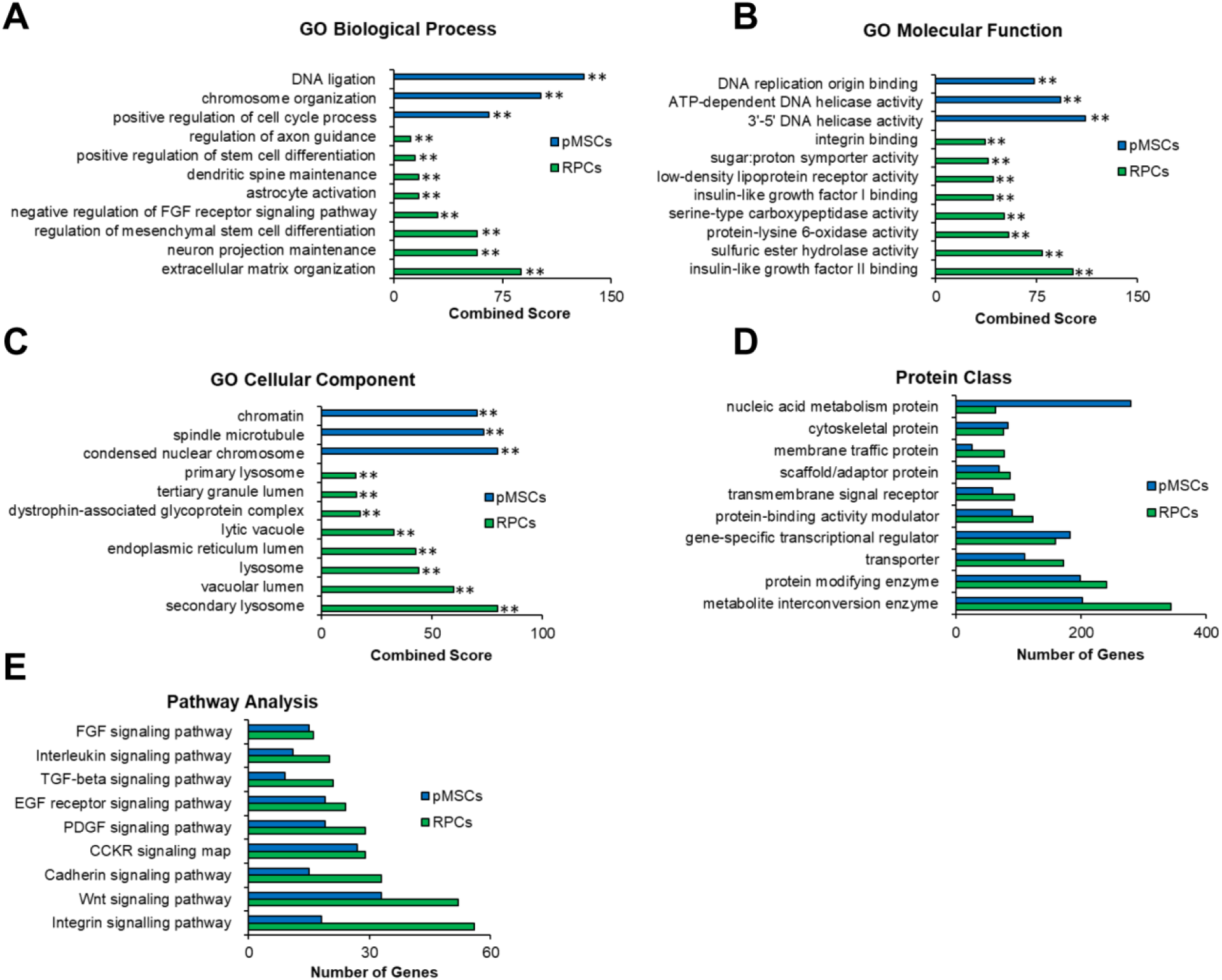
GO analysis of RNASeq of pMSCs and RPCs. Enrichr and PANTHER analyses of DEGs was performed with Benjamini–Hochberg FDR < 0.05 using DESeq2. (A–C) GO term enrichment analysis depicting upregulated DEGs associated with biological processes, molecular function, and cellular component, respectively, and the x-axis represents the combined score generated by Enrichr (**p ≤ 0.01). (D and E) Protein classes and pathways upregulated in pMSCs and RPCs, as determined by PANTHER analysis. The x-axis represents the number of genes associated with each category (**p ≤ 0.01).

Next, we employed PANTHER analysis to investigate differentially expressed protein classes (Fig. 3D), which showed that cytoskeletal and gene-specific transcriptional regulation were enriched in pMSCs and RPCs. However, nucleic acid metabolism proteins were more enriched in pMSCs than RPCs. In contrast, metabolite interconversion enzyme, protein modifying enzyme, transporter, protein-binding activity modulator, transmembrane signal receptor, scaffold/adaptor protein, and membrane traffic protein classes were associated with more genes in RPCs. In signaling pathways, FGF, Interleukin, TGF-beta, EGFR, PDGF, CCKR, Cadherin, Wnt, and Integrin were more expressed in RPCs than pMSCs (Fig. 3E). Altogether, these results showed that RPCs derived pMSCs expressed expected retinal specific genes. Interestingly, they also expressed neurotrophic genes involved in neuroprotection. Therefore, RPCs were an ideal choice to investigate their therapeutic potential.

### Improvement of retinal function in rd12 mice transplanted with cells

Behavioral analysis of rd12 mice intravitreally transplanted with PKH26 labeled pMSCs, and RPCs was performed for vision and acuity using visual recovery assay and optokinetic reflex (OKR), respectively. We observed a significant improvement in visual recovery and acuity in transplanted animals (Supplementary Fig. 1A-C). These observations were followed by a quantitative functional assay using ERG. As expected, ERG analysis showed that dark- and light-adapted rd12 mice had almost a complete loss of a- and b-wave amplitudes (Fig. 4). A progressively significant increase in the a- and b-wave amplitudes of the dark-adapted animals was observed in pMSCs transplanted rd12 mice. Notably, an even greater increase in the a- and b-wave amplitudes was noticed in animals transplanted with RPCs (Fig. 4A-C). The increase in the amplitude was greater at 8 weeks than 4 weeks after cell transplantation, indicating a progressive improvement of the vision in treated mice. In addition, the b-wave amplitude progressively improved with an increase in light intensity in rd12 mice treated with cells, but the increase was more pronounced with RPCs than pMSCs (Fig. 4D-F).

**Figure 4:**
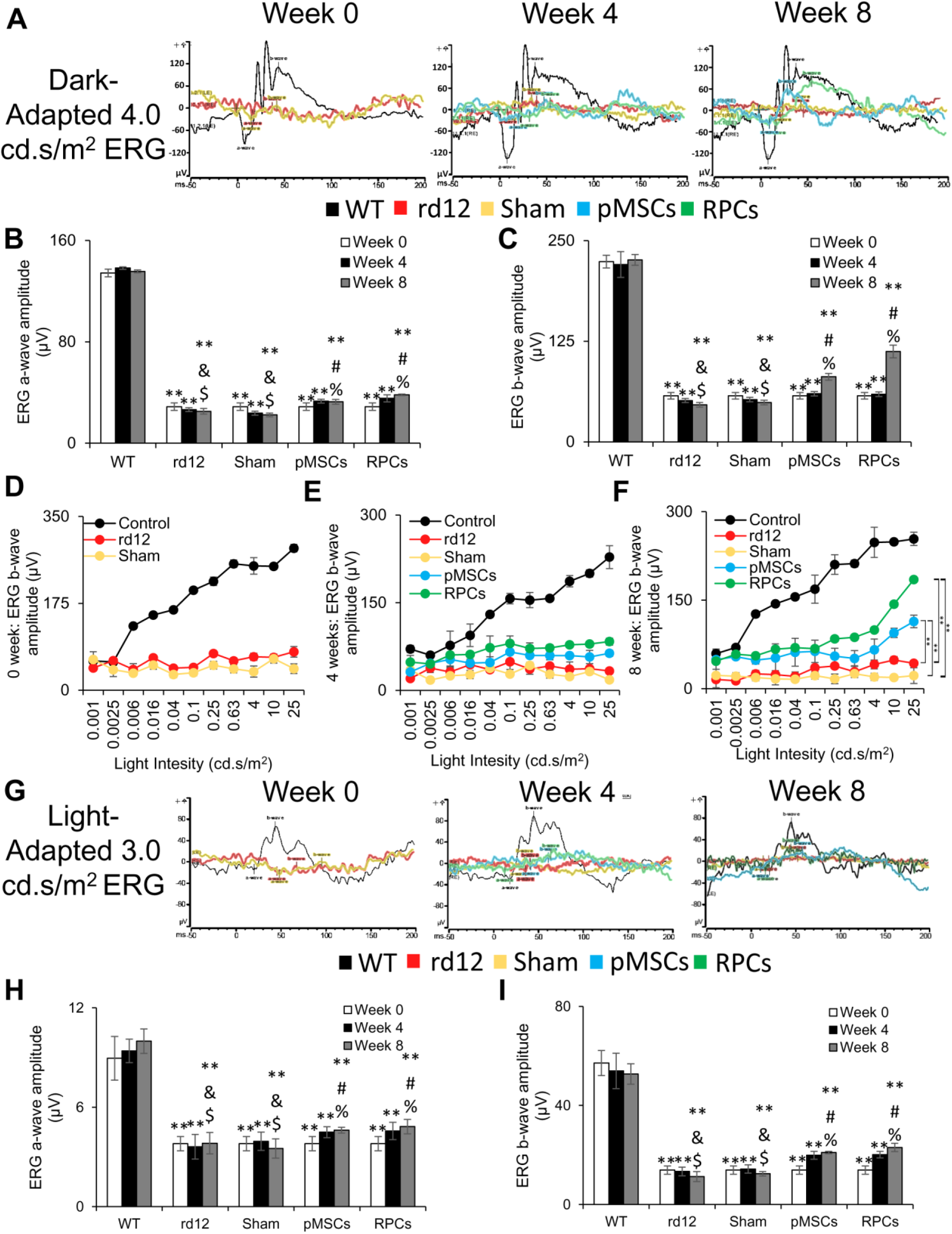
ERG analysis. (A) Representative dark-adapted ERG test of control (wild-type), rd12, sham, rd12+pMSCs and rd12+RPCs at 0, 4, and 8 weeks after transplantation of cells. (B and C) Graphical representation of the amplitude of the dark-adapted ERG a- and b-waves, respectively, from A. (D-F) Intensity-response curves of ERG b-waves at 0, 4, and 8 weeks, respectively, after transplantation of cells (**p < 0.01). (G) Representative light-adapted ERG test of control (wild-type), rd12, sham, rd12+pMSCs and rd12+RPCs at 0, 4, and 8 weeks after transplantation of cells. (H and I) Graphical representation of the amplitude of the light-adapted ERG a- and b-waves, respectively, from G. Symbols, **, #, %, & and $ indicate significant difference at p ≤ 0.01 between all experimental conditions: control (wild-type), rd12, sham, rd12+pMSCs and rd12+RPCs, respectively.

On the other hand, in the light-adapted rd12 mice transplanted with cells, only the b-wave amplitude was progressively increased (Fig. 4G-I). Again, RPCs had a more significant effect in improving the b-wave amplitude than pMSCs, and the effect was more significant at 8 weeks than 4 weeks after transplantation of cells. Overall, these results showed greater improvement in retinal function by RPCs than pMSCs.

### Transplanted cells improved the retinal structure

We next analyzed the structure of the retina. The results depicted in Fig. 5 show that rd12 mice displayed a progressive decrease in thickness of the total retina, ONL, inner nuclear layer (INL), and RGC during the 8 week study signifying degeneration of the retina (Fig. 5A-E). In rd12 mice transplanted with pMSCs, the degeneration of the retina was halted as there was no net decrease in the retinal thickness during the 8 week period. The thickness of the retina, ONL, INL, and RGC significantly increased (from 85.6% to 93.3%, 79.8% to 90.0%, 87.9% to 92.4%, and 80.1% to 86.3%, respectively) by 8 weeks after RPC transplantation. The transplanted cells had no adverse effects as no tumor formation was observed. Overall, pMSCs halted the retinal regeneration while RPCs improved the retinal thickness.

**Figure 5:**
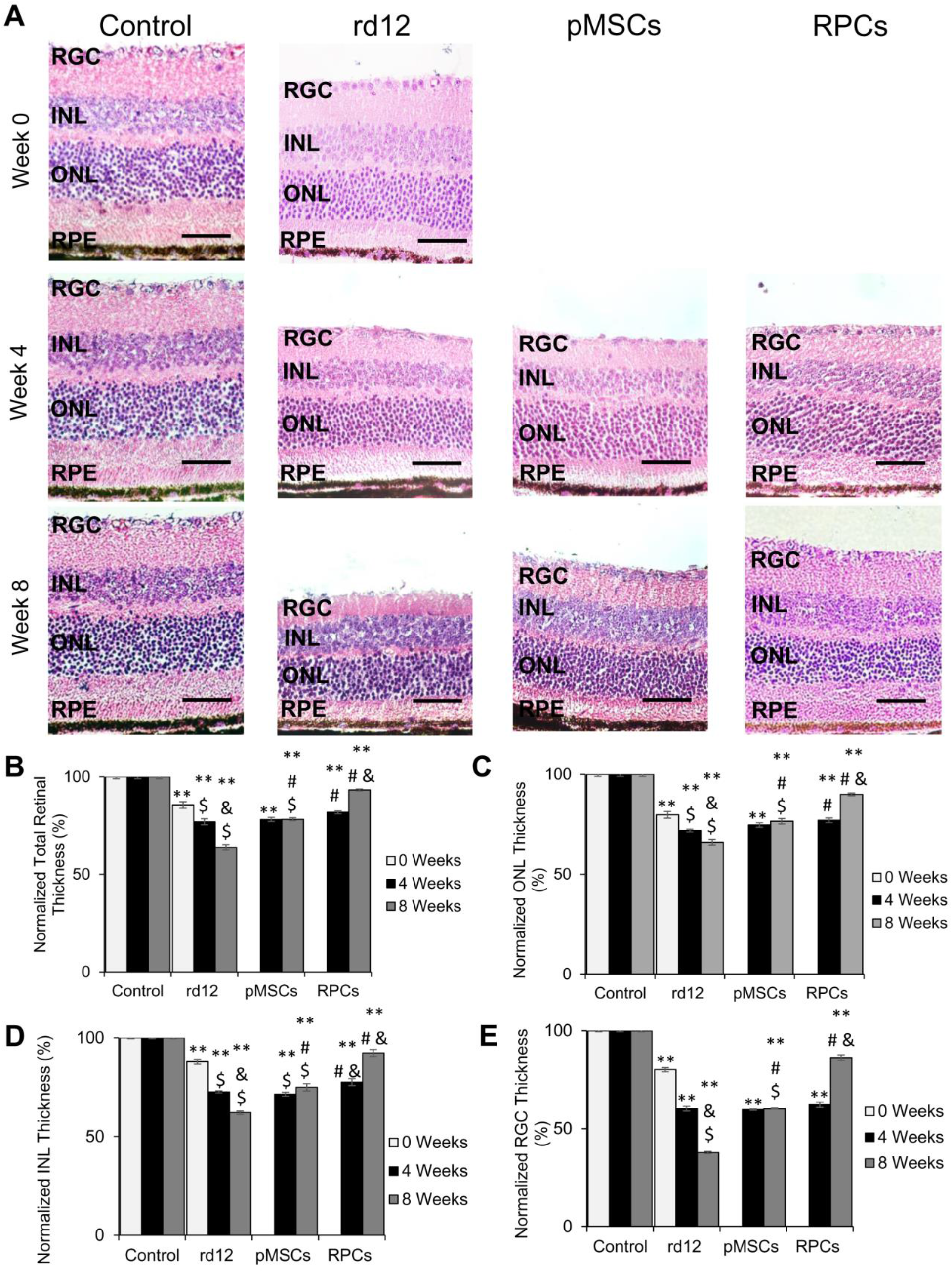
Histological analysis of rd12 retina transplanted with human cells. (A) H&E staining of paraffin-embedded sections of the control (wild-type), rd12, pMSCs and RPCs transplanted retina harvested at 0 week, 4 week, and 8 weeks. All scale bars represent 50 μm. (Magnification: 40x). (B-E) Graphical representation comparing the average thickness of the retina, ONL, INL, and RGC. Thickness of each retina was normalized to the control (wild-type) retina. Symbols, **, #, & and $ indicate significant difference at p ≤ 0.01 between all experimental conditions: control (wild-type), rd12, rd12+pMSCs and rd12+RPCs, respectively.

### Transplanted cells homed to the retina

Cell tracking revealed that PKH26 labeled cells were localized in the retinal whole-mount of animals transplanted with cells (Fig. 6A). Differential interface contrast (DIC) microscopy confirmed the presence and dispersal of labeled cells in the retina (Fig. 6B). RPCs were more dispersed than pMSCs in the retina 8 weeks post-transplantation. These results suggest that transplanted cells homed to different retinal layers.

**Figure 6:**
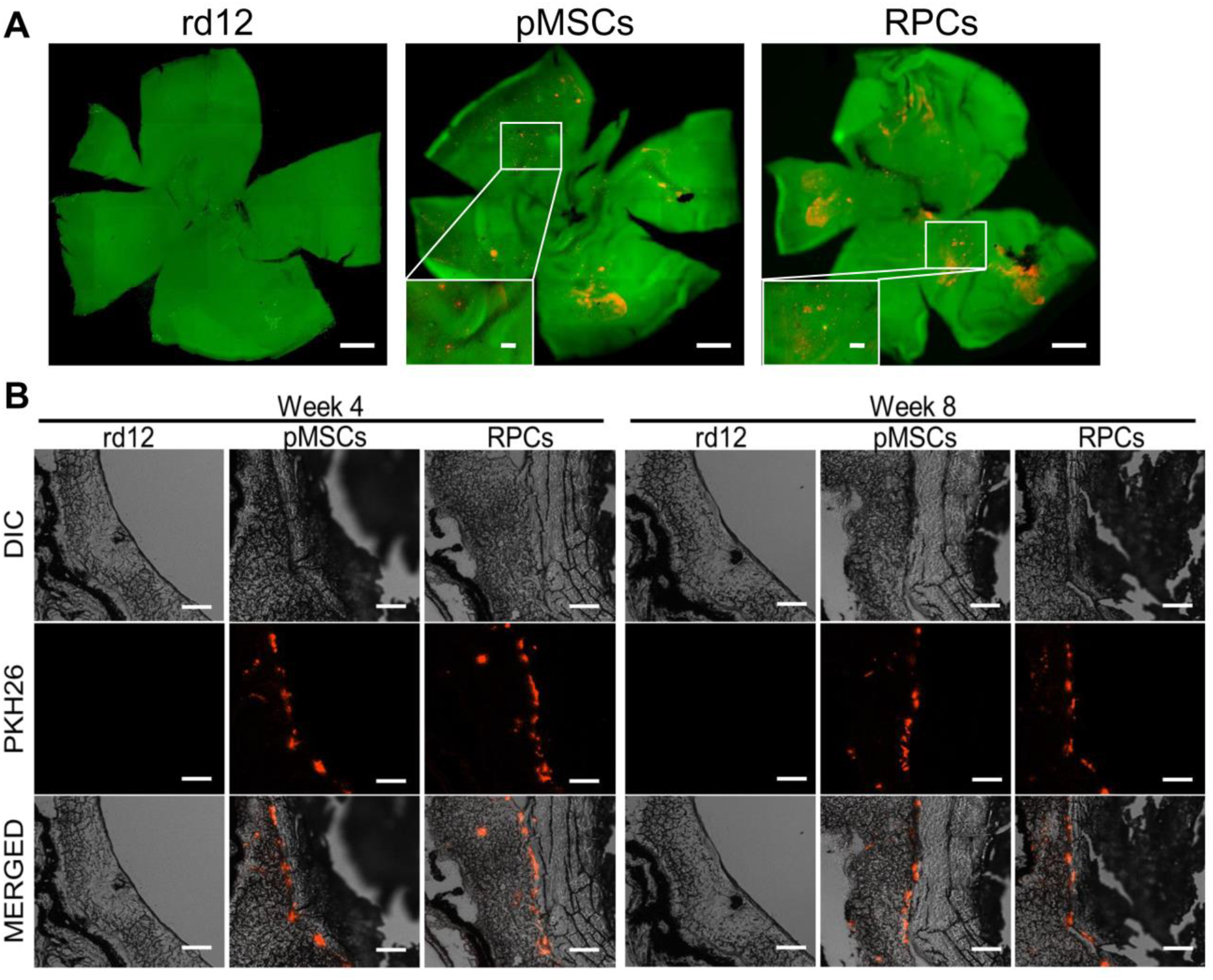
Tracking of cells transplanted into rd12 retina. (A) Whole-mount retina stained with RCVRN (green) 8 weeks after transplantation of PKH26-labeled (red) pMSCs and RPCs. 500 µm scale bar and 20 µm scale bars (inserts). (Magnification: 5x and 40x, respectively). (B) Tracking of PKH26 (red) labeled pMSCs and RPCs in the cryosections of the retina at 4 and 8 weeks. Scale bars represent 100 μm. (Magnification: 20x).

### Expression of human markers in the transplanted rd12 mice retina

Since we found that the transplanted cells were localized in different retinal layers, it was conceivable that they would express various retinal markers. The expression of human markers was first investigated by transcriptional analysis. The results showed the expression of human genes associated with anti-inflammation, neuroprotection, retinal, and neurogenesis. In general, fewer human genes were expressed in animals transplanted with pMSCs than RPCs. Among the anti-inflammatory genes, only expression of *IL-10* was observed 4 weeks after pMSC transplantation, but *IL-4* and *IL-10* were expressed after 8 weeks in the case of RPCs (Fig. 7A). No pro-inflammatory human genes were expressed in either pMSC or RPC transplanted animals.

**Figure 7:**
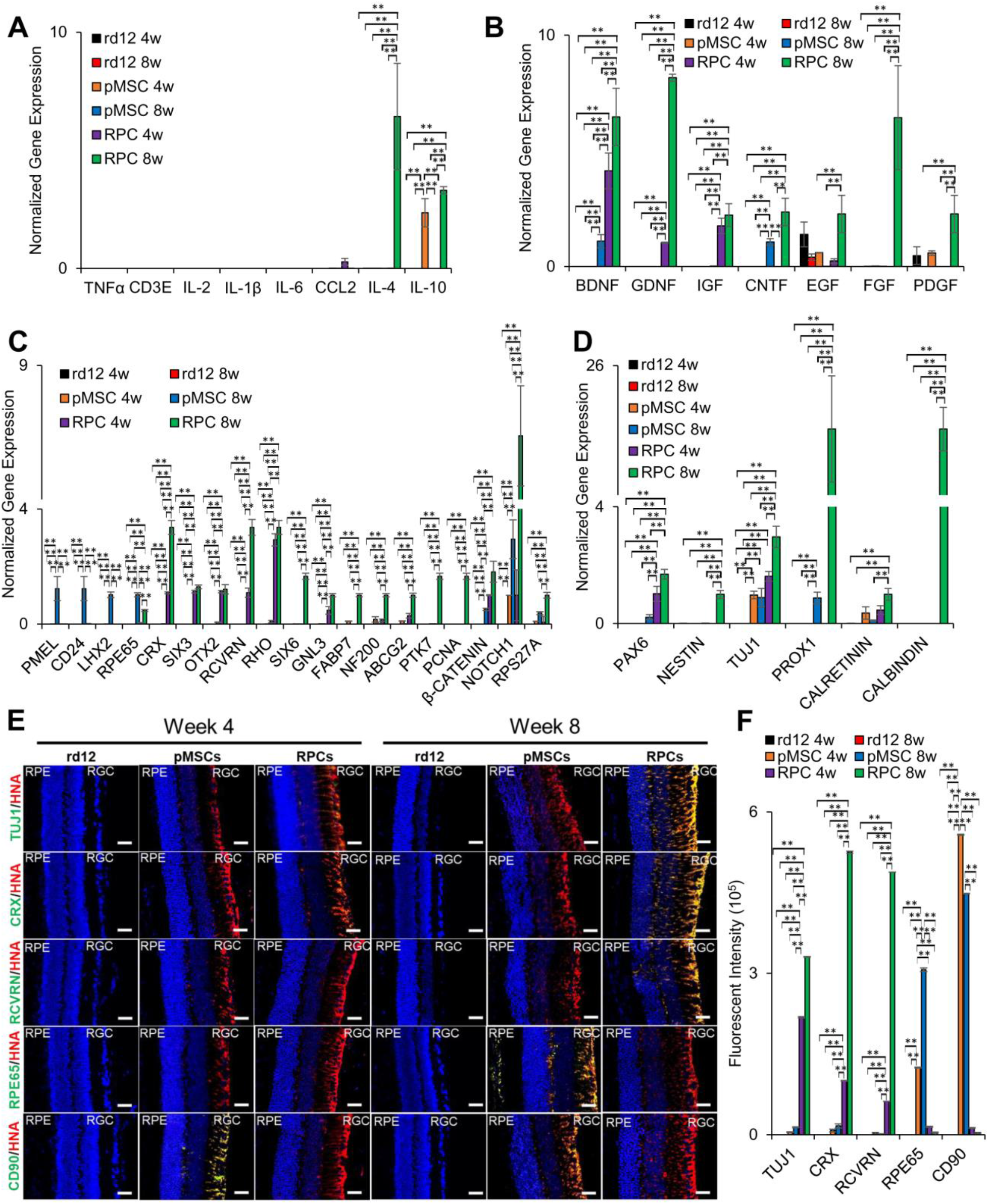
Post-transplantation analysis of the human markers in rd12 retina. (A) Expression of human pro- and anti-inflammatory genes, *TNFα, CD3E, IL-2, IL-1β, IL-6* and *CCL2, and IL-4* and *IL-10,* respectively, (B) neuroprotective genes, *BDNF, GDNF, IGF, CNTF, EGF, FGF, and PDGF,* (C) retinal genes, *CRX, SIX3, OTX2, RCVRN, RHO, SIX6, GNL3, FABP7, NF200, ABCG2, PTK7, PNCA, β-CATENIN, NOTCH1, RPS27A, PMEL, CD24, LHX2,* and *RPE65,* and (D) neurogenesis genes, *PAX6, NESTIN, TUJ1, PROX1, CALRETININ,* and *CALBINDIN*. All gene expression was normalized to *GAPDH* and *β-ACTIN*. Error bars represent the SEM of triplicate experiment (**p ≤ 0.01). (E) Immunohistochemical analysis of paraffin-embedded sections of the retina showing expression of human neural (TUJ1), retinal (CRX, RCVRN, and RPE65), and MSC (CD90) proteins. The sections were countered stained with HNA (red). Blue and green colors represent the DAPI staining of the nuclei and human proteins, respectively. Scale bars represent 50 µm scale bars. (Magnification: 40x). (F) Measurement of fluorescent intensity of proteins as shown in E (**p ≤ 0.01).

The expression of two neuroprotective genes, *BDNF* and *CNTF,* was detected only 8 weeks after transplantation of pMSCs. In RPCs, some neuroprotective genes (*BDNF*, *GDNF*, and *IGF*) were expressed at 4 weeks. However, several genes, *BDNF*, *GDNF*, *IGF*, *CNTF*, *EGF*, *FGF*, and *PDGF* were expressed 8 weeks after transplantation (Fig. 7B).

As for the retinal genes, pMSCs expressed human RPE genes, *PMEL, CD24, LHX2,* and *RPE65*, only at 8 weeks after transplantation. Whereas, RPC transplanted animals expressed human genes, *CRX, SIX3, OTX2, RCVRN*, *RHO, SIX6, GNL3, FABP7, NF200, ABCG2, PTK7, PCNA, β-CATENIN, NOTCH1, and RPS27A* (Fig. 7C), which are predominantly associated with the neural layers (ONL, INL, and RGC) of the retina. The expression of ONL, INL, and RGC genes suggests that transplanted RPCs successfully differentiated and integrated into the neural layers of the retina. In contrast, the expression of RPE genes suggests that pMSCs are differentiated and integrated into the RPE layer of the retina.

Only two neurogenesis genes, *TUJ1* and *PROX1*, were expressed at 8 weeks after transplantation of pMSCs, but several neurogenesis genes, *PAX6, NESTIN, TUJ1, PROX1, CALRETININ*, and *CALBINDIN* were expressed at 8 weeks (with only *PAX6* and *TUJ1* expressed at 4 weeks) after transplantation of RPCs (Fig. 7D).

Similar to the transcriptional analysis, immunostaining studies also suggest that animals transplanted with cells expressed human neural (TUJ1) and retinal (CRX, RCVRN, and RPE65) proteins. These proteins were colocalized with human nuclear antigen (HNA). The results depicted in Fig. 7E-F showed that at 4 weeks after transplantation, HNA-positive cells were found in the ganglion layer and started to migrate towards the INL. However, after 8 weeks of transplantation, HNA-positive cells were more dispersed and further integrated into the retina. We then investigated the expression of cell-specific neural proteins. After 4 weeks of pMSC transplantation, HNA-positive cells expressed CD90 but did not express TUJ1, CRX, RCVRN, or RPE65. However, 8 weeks after pMSC transplantation, most of the HNA-positive cells were found to express RPE65 and were localized in the RPE layer. These results suggest that pMSCs differentiated towards the RPE lineage. In contrast, at 4 weeks after RPC transplantation, almost all HNA-positive cells expressed the neural marker, TUJ1, but a few also expressed the retinal markers, CRX and RCVRN. However, at 8 weeks after RPC transplantation, many HNA-positive cells expressed CRX and RCVRN and were found in the ONL, INL, and RGC layers. RPCs demonstrated differentiation into neuronal lineages. Overall results suggest that transplanted cells differentiated into various retinal cells.

### Transplanted cells promoted endogenous mouse gene expression

To evaluate the effect of the transplanted cells on the rd12 mouse retina, we investigated endogenous mouse genes associated with inflammation, neuroprotection, retina, and neurogenesis. It has been reported that pro-inflammatory genes are upregulated in rd12 mice^[37]^. Our study also found that *Tnfα, Cd3e, Il-2, Il-1β, Il-6,* and *Ccl2* were highly expressed in rd12 mice but not in the wild-type mice (Fig. 8A). However, the expression of these genes was reduced to near wild-type levels in transplanted animals. Interestingly, the expression of anti-inflammatory genes, *ll-4* and *Il-10,* increased in rd12 mice 8 weeks after transplantation of RPCs. Only *ll-10* was expressed at a much lower level in pMSCs than RPCs.

**Figure 8:**
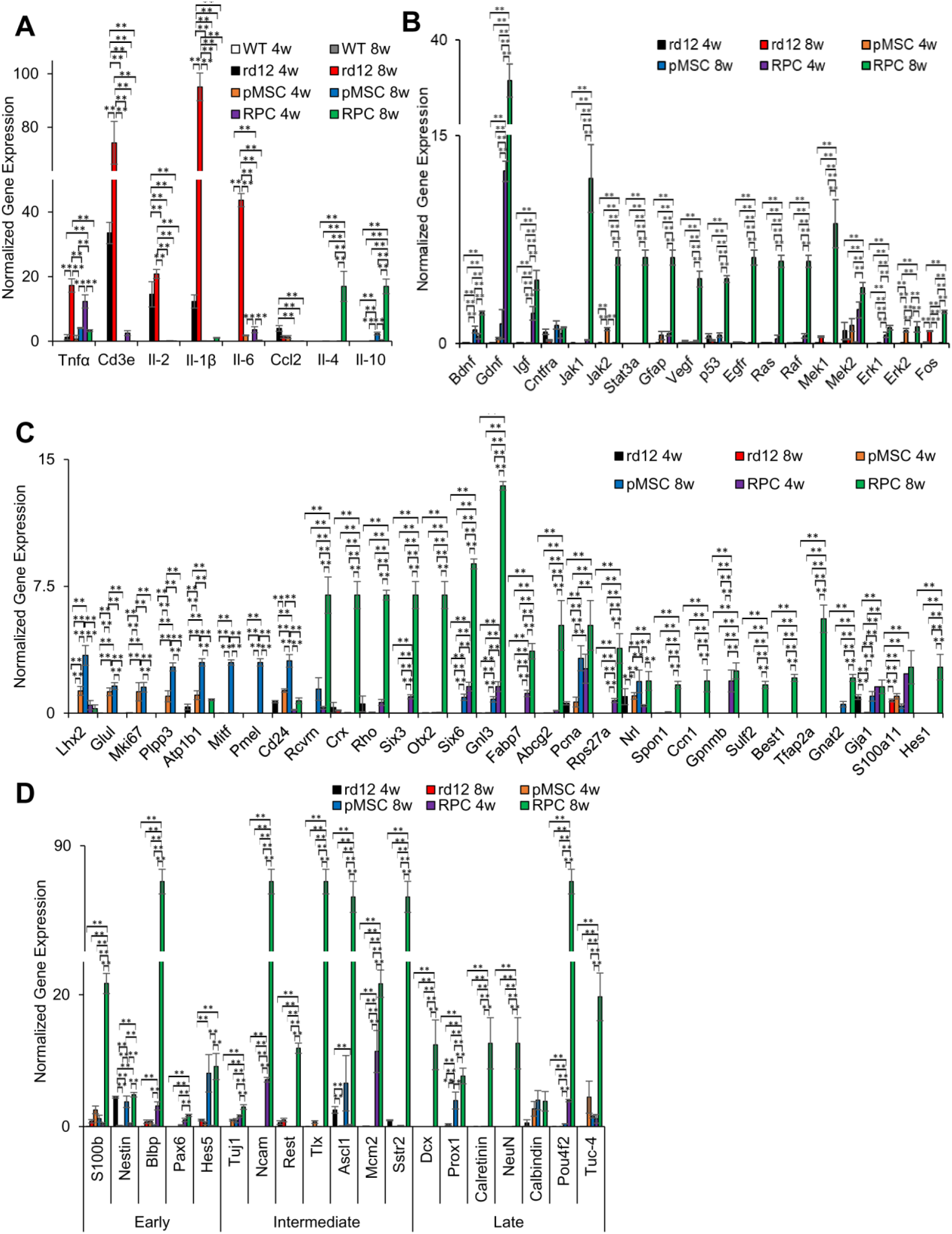
Effect of transplanted cells on the expression of endogenous mouse genes in rd12 retina. (A) Expression of pro-inflammatory genes (*Tnfα, Cd3e, Il-2, Il-1β, Il-6* and *Ccl2) and* anti-inflammatory genes (*Il-4* and *Il-10*),(B) neuroprotective genes *(Bdnf, Gdnf, Igf, Cntfra, Jak1, Jak2, Stat3a, Gfap, Vegf, p53, Egfr, Ras, Raf, Mek1, Mek2, Erk1, Erk2, and Fos*), (C) retinal genes (*Lhx2, Glul, Mki67, Plpp3, Atp1b1, Mitf, Pmel, Cd24, Rcvrn, Crx, Rho, Six3, Otx2, Six6, Gnl3, Fabp7, Abcg2, Pcna, Rps27a, Nrl, Spon1, Ccn1, Gpnmb, Sulf2, Best1, Tfap2a, Gnat2, Gja1, S100Aa11,* and *Hes1) and* (D) genes involved in early (*S100b, Nestin, Blbp, Pax6,* and *Hes5*), intermediate (*Tuj1, Ncam, Rest, Tlx, Ascl1, Mcm2,* and *Sstr2*) and late (*Dcx, Prox1, Calretinin, NeuN, Calbindin, Pou4f2,* and *Tuc-4*) stage of neurogenesis. All gene expression was normalized to *Gapdh* and *β-Actin*, and error bars represent the SEM of the triplicate experiment (**p ≤ 0.01).

Among the neuroprotective genes, *Jak2*, *Erk2*, and *Mek2* as well as *Bdnf* and *Cntfra* were upregulated in the retina of rd12 mice 4 and 8 weeks, respectively, after pMSC transplantation (Fig. 8B). Whereas *Gdnf*, *Igf*, and *Mek2* and *Bdnf, Gdnf, Igf, Jak1, Jak2, Stat3a, Gfap, Vegf, p53, Egfr, Ras, Raf Mek1, Mek2, Erk1, Erk2,* and *Fos* were upregulated in the retina of animals 4 and 8 weeks, respectively, after transplantation of RPCs (Fig. 8B).

In addition, several RPE genes, *Lhx2, Glul, Mki67, Plpp3, Atp1b1,* and *Cd24* were upregulated in the rd12 mice retina 4 weeks after transplantation of pMSCs, and their expression was further increased by 8 weeks. In addition, *Mitf and Pmel* were expressed 8 weeks after transplantation of pMSCs. In the case of RPCs, noticeable expression of *Six3*, *Six6, Gnl3, Fabp7, Pcna, Rps27A, Gpnmb, Gja1, and S100a11* was observed at 4 weeks which further increased by 8 weeks along with expression of several other retinal genes, *Crx, Otx2, Rcvrn*, *Rho, Abcg2, Nrl, Spon1, Ccn1, Sulf2, Best1, Tfap2a,* and *Hes1* (Fig. 8C).

We also saw the upregulation of neurogenesis genes, which are involved in the retina’s early, intermediate, and late stages. For example, the expression of *S100b, Calbindin, and Tuc-4* (after 4 weeks) and *Nestin, Hes5, Ascl1,* and *Prox1* (after 8 weeks) was increased in animals transplanted with pMSCs. Whereas the expression of *Ncam*, *Mcm2*, and *Pou4f2* (after 4 weeks) and *S100b, Nestin, Blbp, Pax6*, *Hes5, Tuj1, Ncam, Rest, Tlx, Ascl1, Mcm2, Sstr2*, *Dcx, Prox1, Calretinin, NeuN, Pou4f2*, and *Tuc-4* (after 8 weeks) was increased in animals transplanted with RPCs (Fig. 8D). Taken together, RPCs were more effective in upregulating the mouse genes involved in inflammation, neuroprotection, retina, and neurogenesis.

### Effect of transplanted cells on the transcriptome of rd12 mouse retina

RNA-seq analysis depicted in Fig. 9A shows hierarchical clustering of rd12 mice vs. rd12 mice transplanted with cells. Expectedly, the untreated rd12 mice formed a clear cluster. In the case of the transplanted animal groups, pMSCs and RPCs showed a higher gene correlation than the untreated animals. Fig. 9B-C identified 39 DEGs (37 upregulated and 2 downregulated genes) at p-value < 0.05 in rd12 mice vs animals transplanted with pMSCs. Whereas 36 DEGs (33 upregulated and 3 downregulated genes) at p-value < 0.05 were found in rd12 mice vs animals transplanted with RPCs (Fig. 9D-E). The comparison between rd12 mice transplanted with pMSCs and RPCs revealed 14 DEGs (11 and 3 upregulated genes in pMSCs and RPCs, respectively) at p-value < 0.05 (Fig. 9F-G).

**Figure 9:**
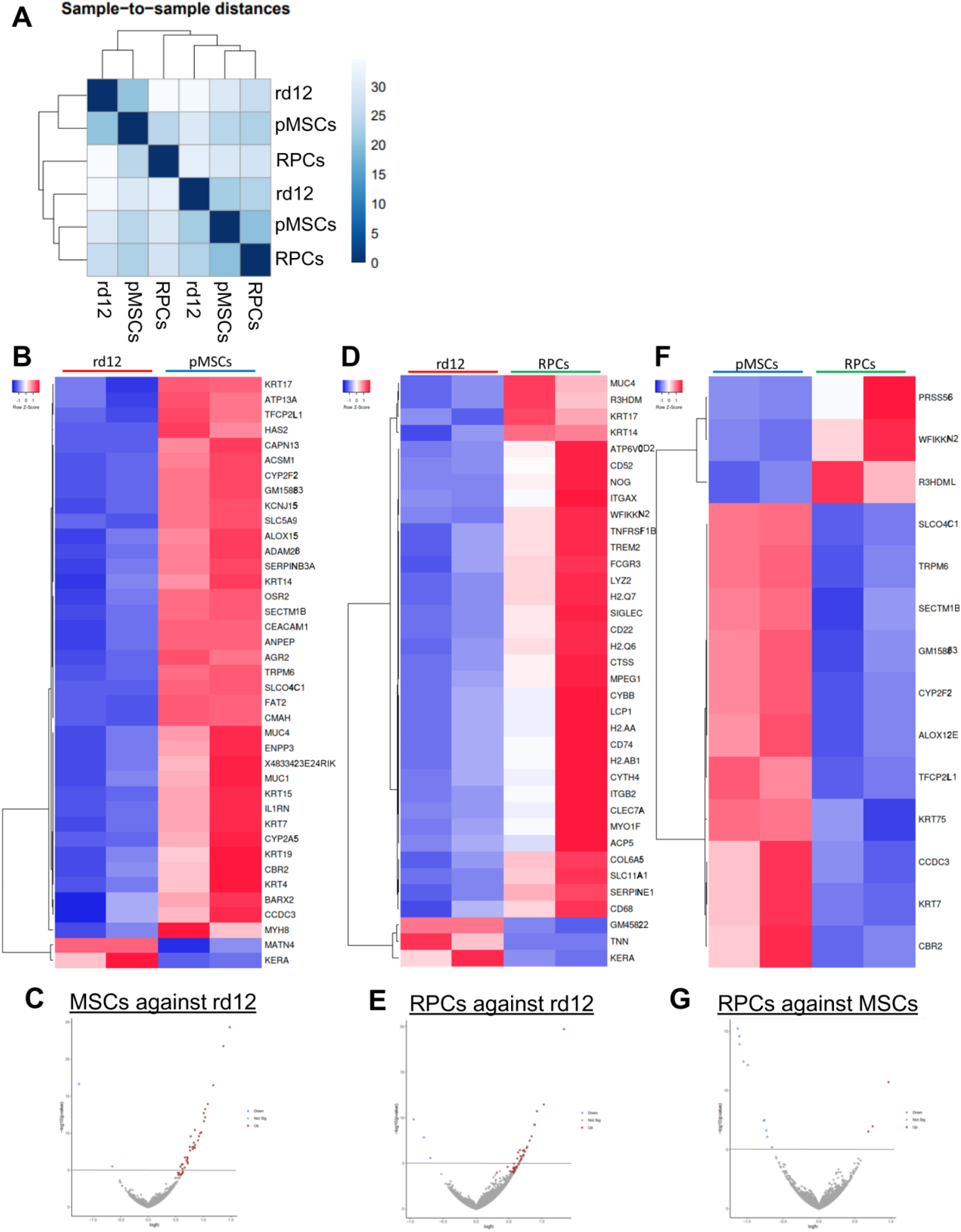
Transcriptome analysis of rd12 mice retina transplanted with cells. (A) RNA-seq data was analyzed to measure sample distances to evaluate for similarities. Hierarchical clustering was performed using DESeq2, and the data was plotted as a sample-to-sample heatmap. The colored indicator represents sample distances (blue signifies a high correlation). (B-G) Heatmap and volcano plots showing raw z-scores of RNA-seq log2 transformed values of the top DEGs in retina of rd12 vs rd12+pMSCs (B-C), rd12 vs rd12+RPCs (D-E), and rd12+pMSCs vs rd12+RPCs (F-G). Upregulated and downregulated genes are red and blue, respectively.

Further analysis of the DEGs, p-value < 0.05 using GO terms identified several GO categories enriched in transplanted animals. In the biological process category, only negative regulation of T cell activation was enriched in rd12 mice transplanted with pMSCs (Fig. 10A). However, in RPC transplanted animals, several processes including response to axon injury, neuron remodeling, regulation of neuroinflammatory response, negative regulation of cytokine production, negative regulation of T cell activation, regulation of cell adhesion mediated by integrin, negative regulation of innate immune response, visual system development, negative regulation of leukocyte activation, positive regulation of glial cell differentiation, neural crest cell migration, regulation of axon extension involved in axon guidance, wound healing, regulation of response to wounding, camera-type eye development, eye development, positive regulation of ERK1 and ERK2 cascade, regulation of ERK1 and ERK2 cascade, and axon guidance were enriched.

**Figure 10:**
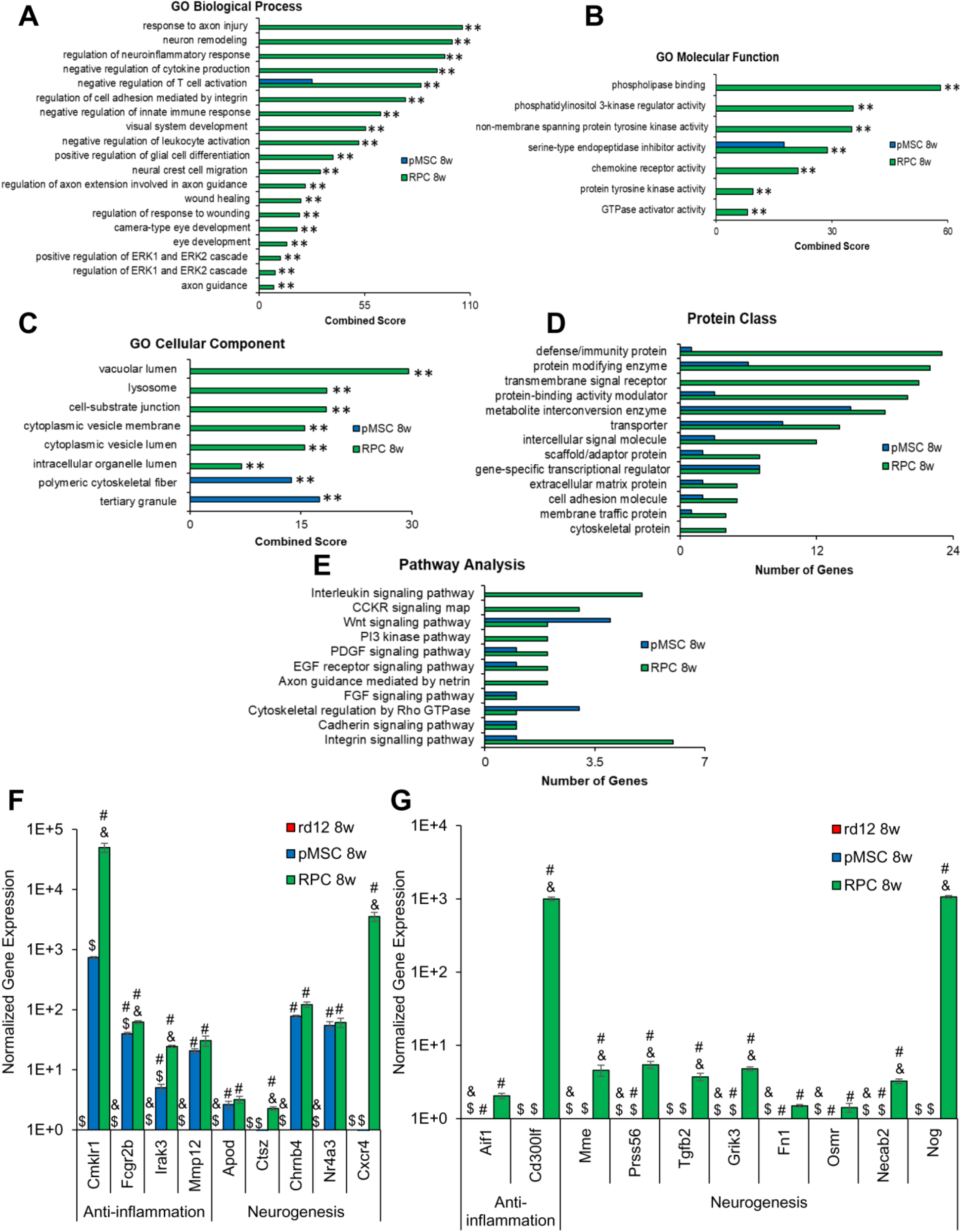
GO analysis of top mouse DEGs in rd12 mice retina transplanted with cells and validation by qRT-PCR. Enrichr and PANTHER analyses of DEGs were performed with p < 0.05 using DESeq2 after 8 week cell transplantation. (A–C) GO term enrichment analysis of upregulated DEGs associated with biological processes, molecular function, and cellular component, respectively, and the x-axis represents the combined score generated by Enrichr (**p ≤ 0.01). (D and E) Protein classes and pathways upregulated in rd12+pMSCs and rd12+RPCs, as determined by PANTHER analysis. The x-axis represents the number of genes associated with each category (**p ≤ 0.01). (F-G) Expression of anti-inflammatory and neurogenesis genes in rd12, rd12+pMSCs and rd12+RPCs and expression ± SEM normalized to rd12 and r12+pMSCs, respectively, as determined by qRT–PCR. Symbols, #, & and $ indicate significant difference at p ≤ 0.01 between all experimental conditions: rd12, rd12+pMSCs and rd12+RPCs, respectively.

In the molecular function category, again, only serine-type endopeptidase inhibitor activity was enriched in rd12 mice transplanted with pMSCs. But several molecular functions including serine-type endopeptidase inhibitor activity, phospholipase binding, phosphatidylinositol 3-kinase regulator activity, non-member spanning protein tyrosine kinase activity, chemokine receptor activity, protein tyrosine kinase activity, and GTPase activator activity were enriched in rd12 mice transplanted with RPCs (Fig. 10B).

In the cellular component category, polymeric cytoskeletal fiber and tertiary granule were enriched in rd12 mice transplanted with pMSCs. Whereas RPCs treated animals were enriched in several cellular components, including vacuolar lumen, lysosome, cell-substrate junction, cytoplasmic vesicle membrane, cytoplasmic vesicle lumen and intracellular organelle lumen (Fig. 10C).

DEGs analyzed by PANTHER revealed that protein classes, metabolite interconversion enzyme, transporter, gene-specific transcriptional regulator, extracellular matrix protein, cell adhesion molecule, and membrane traffic protein were enriched in mice transplanted with pMSCs and RPCs. However, RPC transplanted animals had additional enriched protein classes including defense/immunity protein, protein modifying enzyme, transmembrane signal receptor, protein-binding activity modulator, intercellular signal molecule, scaffold/adaptor protein, and cytoskeleton protein (Fig. 10D).

For pathway analysis, we found that PDGF, EGF receptor, FGF, and cadherin signaling were enriched in rd12 transplanted pMSCs and RPCs. However, Wnt signaling and cytoskeletal regulation by rho GTPase pathways were also enriched in pMSCs. While interleukin, CCKR, PI3 kinase, axon guidance mediated by netrin and integrin signaling pathways were only enriched in the case of RPCs (Fig. 10E).

DEGs were further validated by quantitative gene expression analysis. The results depicted in Fig. 10F-G show that anti-inflammation and neurogenesis genes were significantly upregulated by several log-folds in transplanted animals compared to the rd12 mice. In general, anti-inflammation genes, *Cmklr1, Fcgr2b Irak3* and *Cd300lf,* and neurogenesis genes, *Ctsz, Cxcr4, Mme, Prss56, Tgfb2, Grik3, Necab2,* and *Nog,* were significantly more upregulated in rd12 mice transplanted with RPCs than pMSCs. These results suggest that pMSCs induced genes involved immune response and RPE regeneration. Whereas RPCs promoted expression of genes associated with immune response, neuroprotection, various neural layers of the retina, and neurogenesis.

## Discussion

This study differentiated pMSCs into RPCs to treat retinal degeneration in the rd12 mice, which improved retinal function. Before transplantation, RPCs were characterized for morphological and biochemical properties. pMSC-derived RPCs displayed neural extensions and expressed neural and retinal specific markers similar to published reports^[38–40]^. Transcriptomic analysis revealed that while genes associated with cell growth and proliferation (i.e. *CCNB1, CDC20,* and *CENPF*^[41]^) were highly expressed in pMSCs, they were significantly downregulated in RPCs. In addition, a number of retinal specific genes^[42]^, *GPNMB, BEST1,* and *TYRP1* (RPE), *TFAP2A* and *ATP1B1* (amacrine), *NDRG1* (horizontal), *HES1* and *SPON1* (Müller glia), *SULF2, GNAT2,* and *NRL* (photoreceptor), and *EBF1* (RGC) were upregulated in RPCs. GO analysis of the DEGs indicated that RPCs were enriched in biological processes associated with neuron projection and dendritic spine maintenance involved in neural differentiation^[43]^.

Following cell transplantation, the behavioral analysis showed a significant improvement in visual recovery and acuity of animals. For quantitative analysis of retinal function, the animals were examined by ERG. As expected, both a- and b-waves amplitudes were diminished in untreated animals^[44]^. However, there was a significant improvement in the a- and b-wave amplitudes in the dark-adapted rd12 mice transplanted with cells. Notably, a greater improvement was observed in a- and b-wave amplitudes in animals treated with RPCs than pMSCs. In a previous study, a- and b-wave amplitudes were higher in dark-adapted RCS rats when treated with hBM-MSCs than untreated animals. However, the amplitudes continued to decrease over time^[45]^. Another study reported a combination of hBM-MSCs and fetal RPCs was used to treat retinal degeneration and showed an increase in the a- and b-wave amplitudes of dark-adapted RCS rats compared to untreated animals^[17]^. Again, the amplitudes progressively deteriorated even after cell transplantation. In contrast, subretinal injection of RPE-like cells derived from ESCs into rd12 mice displayed an increase in b-wave amplitude over the untreated animals but demonstrated a diminishing effect over time^[46]^. Feline Müller cells injected into the vitreous of a feline ganglion cell depletion model improved retinal function^[47]^. Unlike these reports, our results for the first time demonstrated a temporal improvement in the a- and b-wave amplitudes, particularly in the case of RPCs, suggesting lasting effect of cell transplantation.

Post-transplantation histological analysis revealed temporal degeneration of retina in rd12 mice as previously described^[48]^. The overall retinal thickness in rd12 mice was 86%, 77%, and 64% compared to the wild-type animals at 0, 4, and 8 weeks, respectively. However, the thickness was 78% at both 4 and 8 weeks in pMSC treated animals and 82% and 93% at 4 and 8 weeks, respectively, in the case of RPCs. These results suggest that while pMSCs provided protection that led to retinal thickness maintenance, RPCs halted degeneration and promoted retinal regeneration. In general, the thickness of all three nuclear layers, ONL, INL, and RGC, was significantly and progressively increased in the cell transplanted animals compared with rd12 mice. However, more improvement was observed in the ONL, INL, and RGC in RPC transplanted mice. In reported studies, transplanted MSCs have improved either ONL or RGC but not both^[12, 49]^. Therefore, our results are significant and exciting as they exhibited improvement in almost all nuclear layers, particularly in RPCs.

Previously, ESC retinal derivatives were found to survive and migrate between the ONL and RPE layers upon subretinal transplantation^[46]^. Whereas fetal RPCs were found to migrate towards the INL and ONL^[34]^. However, our cell tracking analysis showed localization of labeled cells at various sites of the retinal whole-mount, suggesting that transplanted cells dispersed from the injection site. DIC analysis also indicated the localization of labeled cells in multiple layers of the retina.

Since, MSCs are known to exert paracrine effects^[50]^, we investigated the expression of human inflammatory cytokines and neurotrophic factors in the retina of rd12 mice. Our results indicated high levels of xenogenic expression of human anti-inflammatory and neuroprotective genes in animals treated with pMSCs and RPCs. No pro-inflammatory markers were expressed, suggesting that transplanted cells did not exert an immune response and were safe for cell therapy. Several human retinal and neurogenesis genes were expressed in the transplanted animals, presumably providing neural protection and regeneration. Supporting these findings, xenogeneic protein expression of retinal markers was also observed in the retina of transplanted animals. In general, xenogeneic expression was more significant in RPC than in pMSC transplanted animals. Although several reports studied the effectiveness of cells in treating RDD in animal models^[45, 49, 51]^, xenogenic expression of neuroprotective and neurogenesis markers has not been well-investigated. It is conceivable that retinogenesis may in part be due to the differentiation of transplanted cells.

In retinal degeneration, expression of pro-inflammatory genes has been reported to increase with a subsequent decrease in the expression of retinal genes and vision loss^[11, 13, 52]^. We also found upregulation of several mouse pro-inflammatory genes, *Tnfα, Cd3e, Il-2, Il-1b, Il-6,* and *Ccl2*, and downregulation of several retinal genes in rd12 mice. Therefore, we investigated the effect of transplanted cells on the expression of inflammatory and retinal genes. Our study showed that transplanted cells induced endogenous inflammatory response by suppressing pro-inflammatory genes while activating anti-inflammatory and retinal genes. RPCs were more potent than pMSCs in exerting these beneficial effects. We also found that the transplanted cells, specifically RPCs, significantly upregulated several endogenous neurotrophic factors important for cell proliferation, survival, and neuronal differentiation^[53, 54]^. This observation supports the results determined improvements in OKR, ERG, and retinal structure.

In contrast to birds, amphibians, and fish, *de novo* neurogenesis and regenerative capacity of the adult mammalian retina is very limited^[55, 56]^. Therefore, only a few studies have investigated neurogenesis resulting from the augmentation of cells into the mammalian retina^[9, 57]^. We observed that mouse RPE and neural gene expression was increased in animals transplanted with pMSCs and RPCs, respectively. In addition, we investigated neurogenesis genes in the retina of animals transplanted with cells. The expression of only a few genes, *Nestin, Hes5,* and *Prox1,* increased 8 weeks after transplantation of pMSCs. Whereas the expression of several genes involved in early, intermediate, and late neurogenesis was significantly increased in animals transplanted with RPCs, and their expression was even greater 8 weeks after transplantation. Altogether, our results suggest that pMSCs may be involved in the regeneration of mouse RPE. Whereas RPCs promoted regeneration of the ONL, INL, and RGC. They also validate the improvement in retinal structure and function, as stated earlier. Nevertheless, further studies would be helpful to determine the specific cell types involved in the regeneration of rd12 mice retina.

The transcriptomic analysis also revealed significant differences in the animals treated with cells compared to rd12 mice. Analysis of RNA-seq data showed that large dispersions in the sample groups did not affect differential gene expression. We found a clear and significant difference in the DEGs by analyzing the gene expression pattern affected by pMSCs and RPCs. Comparative heatmap and volcano plot analyses revealed differences between animals transplanted with pMSCs and RPCs only in a select number of DEGs. GO analysis indicated several biological processes enriched in the transplanted animals. In the case of pMSCs, DEGs were involved in the suppression of T cell activation. Whereas RPCs not only suppressed the expression of genes involved in the immune response but also promoted the enrichment of biological processes associated with neuron remodeling, visual system, and eye development, as well as axon guidance. This pattern of enrichment suggested that the transplantation of pMSCs and RPCs were effective in promoting anti-inflammatory response. RPCs promoted retinal regeneration, which led to greater improvement in visual function. These findings complement the ERG and visual recovery results as discussed earlier.

When DEGs were subjected to PANTHER analysis, activation of several pathways in the transplanted animals were revealed. Most of the signaling pathways involved in neural development were activated in both pMSCs and RPCs, except that interleukin, CCKR, PI3k, and axon guidance signaling pathways were activated only in RPCs. qRT-PCR analysis of DEGs validated the activation of genes involved in anti-inflammation and neurogenesis, which were expressed at higher levels in RPCs than pMSCs. Since qRT-PCR is a highly sensitive technique, the differences in DEG expression were many log fold higher. Some highly expressed genes in RPC transplanted animals were also associated with anti-inflammatory response^[58]^. While the other highly expressed genes are known to play a role in neuronal and retinal development. For example, *Necab2* and *Prss56*, are associated with neuronal development and ocular axial growth, respectively^[59, 60]^. Another gene, *Grik2* acts as an excitatory neurotransmitter^[61]^. Altogether, these results demonstrated that RPCs countered inflammation and induced neuroprotection and neurogenesis that improved retinal structure and physiological function.

The molecular analysis of RPC transplanted animals led us to propose a mechanism for improving retinal protection and regeneration (Supplementary Fig. 2). In this mechanism, upregulation of CNTF could activate the JAK/STAT pathway. RPC transplanted animals expressed high levels of CCN1 that can also activate this pathway. In addition, activated STAT3α can turn on the genes (i.e., *Gfap, Vegf* and *p53*) involved in cell proliferation and survival. In addition, EGF/FGF may activate MAPK signaling pathways, which turn on the *Ccnd1*gene, known to be involved in proliferation and neuronal differentiation. Lastly, NOG will bind to the BMPR to inhibit the BMP pathway, which can also be inhibited by FGF signaling.

In conclusion, our study demonstrated that RPCs counter inflammation, provided retinal protection, and promoted neurogenesis, resulting in improved retinal structure and physiological function in rd12 mice. Although several small clinical trials have utilized BM-MSCs and UC-MSCs, the results are not always conclusive^[27, 62, 63]^. Our investigation provided proof of concept and basis for large-scale preclinical and clinical studies to treat RDD using RPCs.

## Materials and Methods

### Maintenance and culture of pMSCs

Previously isolated, well-characterized, and highly proliferative pMSCs, were maintained using a growth medium containing DMEM nutrient mix F12 medium (DMEM/F12; Life Technologies, Carlsbad, CA, USA), supplemented with 10% fetal bovine serum (FBS; VWR, Radnor, PA, USA), and 5.6% of antibiotic solution (0.1% gentamicin, 0.2% streptomycin, and 0.12% penicillin); (Sigma, St Louis, MO, USA) and incubated at 37°C in an atmosphere of 5% CO_2_ in a humidified incubator.

### Differentiation of pMSCs into RPCs

pMSCs were induced to differentiate towards the retinal lineage by culturing them in neurobasal media (Thermo Fisher Scientific, Waltham, MA, USA) containing 50 ng epidermal growth factor (EGF; PeproTech, Rocky Hill, NJ, USA), 1 µM retinoic acid (Sigma), 100 µM taurine (PeproTech), 2 mM glutamine (Sigma), 1x B27 supplement (Thermo Fisher Scientific), and 2% FBS for 2 weeks. The differentiated cells were then characterized by flow cytometry, qRT-PCR, and immunocytochemical staining analyses.

### Flow cytometric analysis

Cells were grown to 70% confluency and stained against CD44 and CD90 (FITC labeled antibodies) or CD29, CD73, CD105, and RCVRN (APC labeled antibodies) (Becton Dickinson, Franklin Lakes, NJ, USA; Santa Cruz, Dallas, Tx, USA, respectively). Labeled cells were analyzed using FACS Canto II (Becton Dickinson) and Diva Software (Becton Dickinson). APC- and FITC-labeled mouse IgG were used as negative controls.

### Immunocytochemical analysis

Cells were fixed with 4% paraformaldehyde for 10 minutes at room temperature, permeabilized with 0.5% Triton X-100 (Sigma) and blocked with 2% bovine serum albumin (Sigma) for 1 hour. Cells were then stained with primary antibodies at 1:100 dilution at 4°C overnight, followed by staining with secondary antibodies at 1:200 dilution at room temperature for 2 hours. Cells were counterstained with DAPI at 1:100 dilution for 5 minutes at room temperature. Fluorescent images were captured using a confocal microscope (NIKON Instruments Inc, Melville, NY, USA). Fluorescent intensity was calculated using ImageJ software (NIH, Bethesda, MD, USA).

### Animal studies

All animal experiments were approved by the Institutional Animal Care and Use Committee (IACUC), Providence Hospital, Southfield, Michigan (IACUC #101-16) and Oakland University, Rochester, Michigan (IACUC #19082), as well as the Institutional Biosafety Committee (IBC), Oakland University, Rochester, Michigan (IBC #2858). A total of 84 rd12 mutant mice from Jackson Laboratory (Bar Harbor, ME, USA) were used for this study. The rd12 mice were set up in breeding triads with one male and two females and received 7-9 pups per litter. C57BL/6J mice were used as controls. All mice were maintained at the Providence Hospital Animal Facility and Oakland University Animal Facility under a 12/12-hour light and dark cycle.

### Cell transplantation in an rd12 mouse model

Cells were labeled with cell membrane labeling dye PKH26 (Sigma) following the manufacturer’s instructions. Confocal microscopy was used to confirm the efficiency of the fluorescently labeled cells prior to transplantation. Six week old rd12 mice were anesthetized by intraperitoneally injecting 0.1mL/10g mixture of ketamine/xylazine (ketamine (50 mg/kg) and xylazine (7 mg/kg)). A drop of topical anesthesia (Proparacaine Hydrochloride, Ophthalmic Solution, Alcaine, Acon, Canada) was applied to the eye before transplantation, and pupils were dilated with 1% atropine and 2.5% phenylephrine hydrochloride (Ophthalmic Solution). Using a 27 G sharp disposal needle, the eye was punctured to create a hole by a blunt needle. The animals were intravitreally injected with 1 µL of DMEM containing PKH26 labeled pMSCs (2 x10^5^) or RPCs (2 x10^5^) through a 32 G needle, using a 10 µL Hamilton syringe. The experiments were performed in triplicates. The animals were monitored regularly for 4 or 8 weeks after cell transplantation.

### Visual function analysis

All visual function analyses were performed the day of transplantation and every two weeks following cell transplantation. To measure visual recovery, mice were placed in a small cage for 5 minutes before testing. A long Q-tip was then placed in front of the eye, and the reaction time of the mouse moving its head away or recognizing the Q-tip was recorded. A score from 0-4 was given to each test (0: No head movement, 1: >1 second delay for head movement, 2: 1 second delay for head movement. 3: <1 second delay for head movement, and 4: Immediate head movement). The non-transplanted rd12 mice were used as negative controls. The C57BL/6J mice were used as positive controls. Visual recovery scale is found in Supplementary Fig. 1A. For the visual acuity, optokinetic response (OKR) was analyzed. The mice were placed onto a small cylinder in the center of the apparatus to allow full movement. Monitors were attached to the walls of the apparatus. A video that displayed vertical alternating black and white lines moved from left to right of the screen at a constant speed. The movement of the mouse was recorded. To obtain the visual acuity of each condition, the cycles per degree (cpd= 1/[2arctan(h/2a)]) was calculated. The non-transplanted rd12 mice served as negative controls. The C57BL/6J mice were used as positive controls.

### Electroretinography

To investigate the therapeutic effect of transplanted cells on the retina function, ERG analysis was performed 1 week before the transplantation of cells in all mice and every other week thereafter. Full-field ERGs were recorded with an Espion-III system (Diagnosys LLC, Lowell, MA, USA) connected to a computer-based system. ERGs of age-matched non-transplanted rd12 eyes were used as negative controls, and C57BL/6J mice were used as positive controls. All testing was performed in a climate-controlled, electrically isolated dark room under dim red light illumination. Mice were dark-adapted for 4 hours and anesthetized with ketamine (50 mg/kg) and xylazine (7 mg/kg). Body temperature was maintained on a 37°C warming pad. Proparacaine hydrochloride droplets were used to anesthetize the eyes, and pupils were dilated with 1% atropine and 2.5% phenylephrine hydrochloride. Methylcellulose gel (Ophthalmic Solution) was applied to the eyes, and the wire loop electrodes were placed on the cornea. Needle reference and ground electrodes were inserted into the cheek and tail, respectively. Dark-adapted mice were tested with a series of dim light flashes (0.001-25 cd.s/m^2^). Eyes were then light-adapted to a constant background light for 10 minutes. Then the ERG response traces were averaged from a series of rapidly flickering bright light flashes (15 to 30 flashes per second at an intensity of 3 cd.s/m^2^) over several seconds.

### Tracking of transplanted cells and histological analysis

The animals were sacrificed by CO_2_ overdose, and the eyes were harvested 4 weeks or 8 weeks post-transplantation. For paraffin embedding, eyes were fixed in 97% methanol and 3% acetic acid for 48 hours at −80°C. Eyes were then transferred to −20°C for 4 hours and then 48 hours at room temperature. Eyes were then dehydrated in 100% ethanol for 15 minutes and repeated thrice, and then transferred to xylene for 15 minutes and repeated twice. Eyes were then embedded in paraffin and sectioned (5-10 μm thick) using a microtome. The paraffin-embedded retina was stained with hematoxylin and eosin (H&E) (Thermo Fisher Scientific) to evaluate the cellular structure of the retina. To detect the protein levels of the human retina specific markers within the mouse retina, the sections were stained with primary antibodies 1:100 dilution at 4°C overnight followed by staining of secondary antibodies at 1:200 dilution at room temperature for 2 hours. Cells were counterstained with DAPI at 1:100 dilution for 5 minutes at room temperature. Fluorescent images were captured using a confocal microscope (NIKON Instruments Inc.).

For OCT embedding, eyes were fixed in 4% paraformaldehyde for 10 minutes, placed in 10% sucrose for 30 minutes, and then in 30% sucrose overnight at 4°C. The eyes were then embedded in OCT medium (Thermo Fisher Scientific) and sectioned (6–10 μm thick) using a cryostat. The sections were observed under confocal microscopy for presences of PKH26 dye (Cy3) labeled cells throughout the retina. To mount for cell tracking, the retinas were flattened with 4-5 relaxing cuts, fixed in 4% paraformaldehyde, stained with anti-mouse RCVRN primary antibody, followed by staining of Alexa flour 488 secondary antibodies (Thermo Fisher Scientific) and analyzed by fluorescent images captured using a confocal microscope (NIKON Instruments Inc.).

### qRT-PCR analysis

Isolation of the total cellular mRNA from cells and tissue was done using the GeneJET RNA purification kit (Thermo Fisher Scientific) following the manufacturer’s instructions. Total RNA was purified with DNase and incubated at 37°C for 30 minutes using a thermocycler (Bio-Rad, Hercules, CA, USA). cDNA was synthesized using an iScript kit (Bio-Rad). qRT-PCR was performed by using Sso-Advanced Universal SYBR Green Supermix Kit (Bio-Rad) on CFX96 Real-Time System (Bio-Rad). A 10 µL reaction was used, which included 5 µL Sybr green, 3 µL of distilled water, 0.5 µL of forward primer, 0.5 µL of reverse primer, and 1 µL of 1:10 diluted cDNA. Each reaction was exposed to the following conditions: 98°C for 10 minutes, followed by 30 seconds of 98°C, 20 seconds of 60°C, and 30 seconds of 72°C for 44 cycles in 96-well optical reaction plates (Bio-Rad). The reference genes, GAPDH, and β-ACTIN, were used to normalize the targeted genes. Each reaction was performed in triplicate. Primers were screened using primer BLAST^[64]^ for homology. Human and mouse-specific primer sequences are listed in Supplementary Table 1 and Supplementary Table 2.

### Analysis of retina thickness

The retina embedded in paraffin was sectioned at 5-10 µm and stained with H&E. The same location of the retina was selected for each eye, and the thickness of the whole retina, ONL, INL, and RGC was measured by Image J (NIH, Bethesda, MD, USA).

### RNA-sequencing analysis

RNA-seq analysis was performed as previously described^[65]^. In brief, the total RNA of the samples was isolated using the GeneJet RNA purification kit (Thermo Fisher Scientific). cDNA library was prepared for RNA-seq using KAPA RNA HyperPrep Kit with RiboErase (Kapa Biosystems Inc.). The cDNA libraries were sequenced using Illumina HiSeq machine for paired-end reads using GENEWIZ (South Plainfield, NJ) services, and sequencing analysis was performed on the GALAXY platform^[66]^. Raw sequencing reads were aligned to the human genome assembly hg38, and reads were counted and annotated using featureCounts using GENCODE GRCh38 comprehensive gene annotation. DESeq2 was used to normalize counts and determine DEGs using statistical analysis. Ontology enrichment analysis of DEGs was performed using PANTHER and Enrichr software^[67, 68]^.

### Statistical analysis

Data are presented as the mean ± standard error of the mean (SEM) of triplicates per analysis. Results with **p ≤ 0.01 were considered statistically significant. All analyses were performed using SPSS version 26 (SPSS Inc. USA) using the one-way ANOVA test.

## Data availability

The raw and processed sequencing data generated in this study have been deposited in the NCBI GenBank database and is pending for the accession code.

## Acknowledgments

We like to thank Dr. Dao-Qi Zhang and Dr. Tomomi Ichinose for help with the ERG analysis. We also like to thank Dr. Chhabi Govind for reviewing the manuscript. Dr. R. Dodd, Jessica Lenyard and Janet Schofding for assistance in animal studies at Providence Hospital and Oakland University. The study was supported by the OU-WB Institute for Stem Cell and Regenerative Medicine (ISCRM), Oakland University, Rochester, MI, Ascension – Providence Hospital, Southfield, MI, and Michigan Head and Spine Institute, Southfield, MI. We also acknowledge support from Oakland University for the Provost Graduate (C. Brown and C. McKee) and Undergraduate (K. Walker and M. Mazzella) Research Awards.

## Author information

### Contributions

Conceptualization: G.R.C., C.B; Methodology: G.R.C., C.B., K.W., and P.A.; Validation: C.B., K.W., C.M., and G.R.C.; Formal Analysis: C.B., K.W., M.M., and G.R.C.; Investigation: G.R.C., C.B., K.W., P.A., and D. S.; Resources: G.R.C.; Data curation: C.B., and G.R.C.; Supervision: G.R.C.; Visualization: C.B., K.W., and G.R.C.; Funding acquisition: G.R.C.; Project administration: G.R.C.; Writing – original draft: C.B. and G.R.C.; Writing – review & editing: all authors. All authors have read and approved the manuscript.

---

**Supplementary Table 1.**
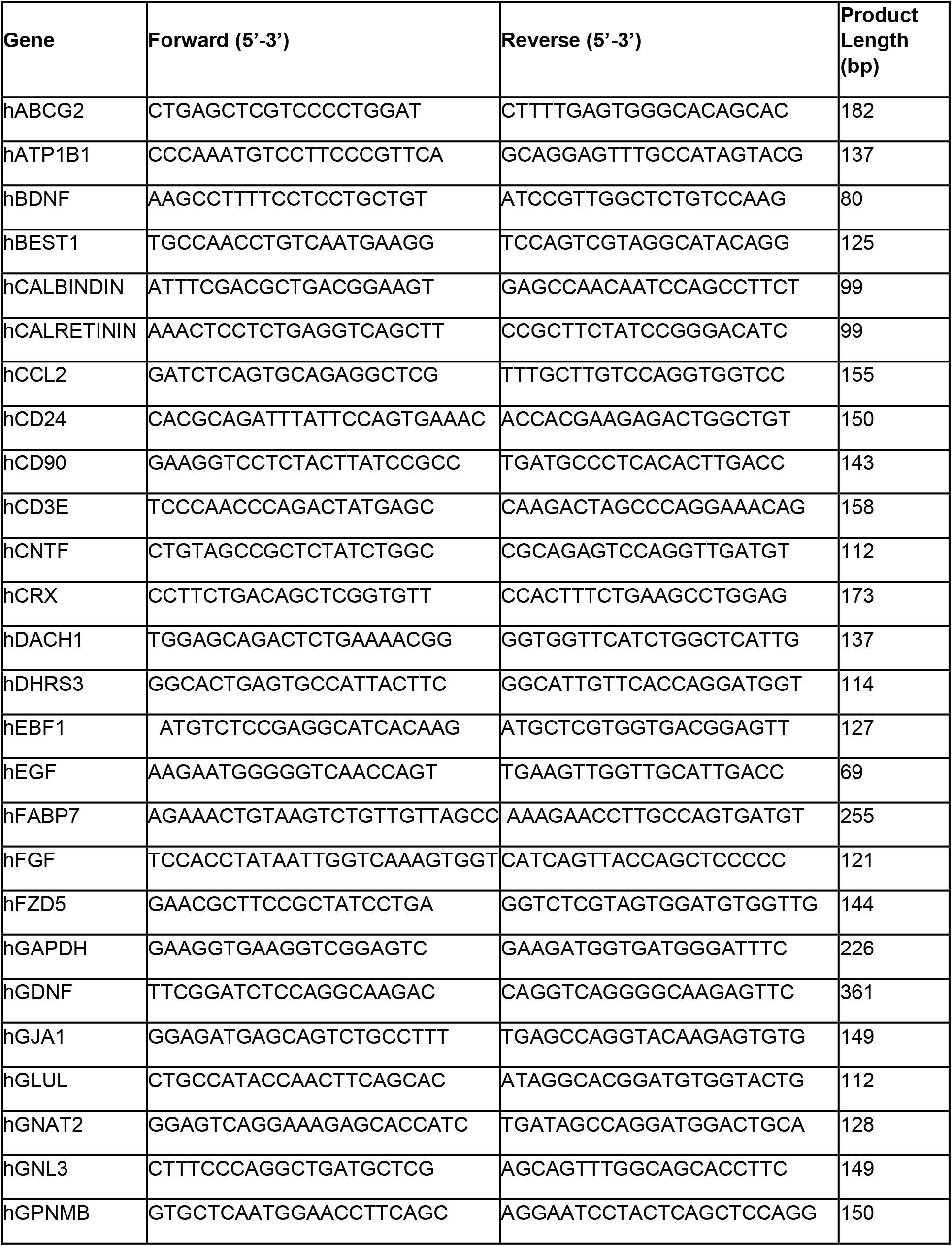
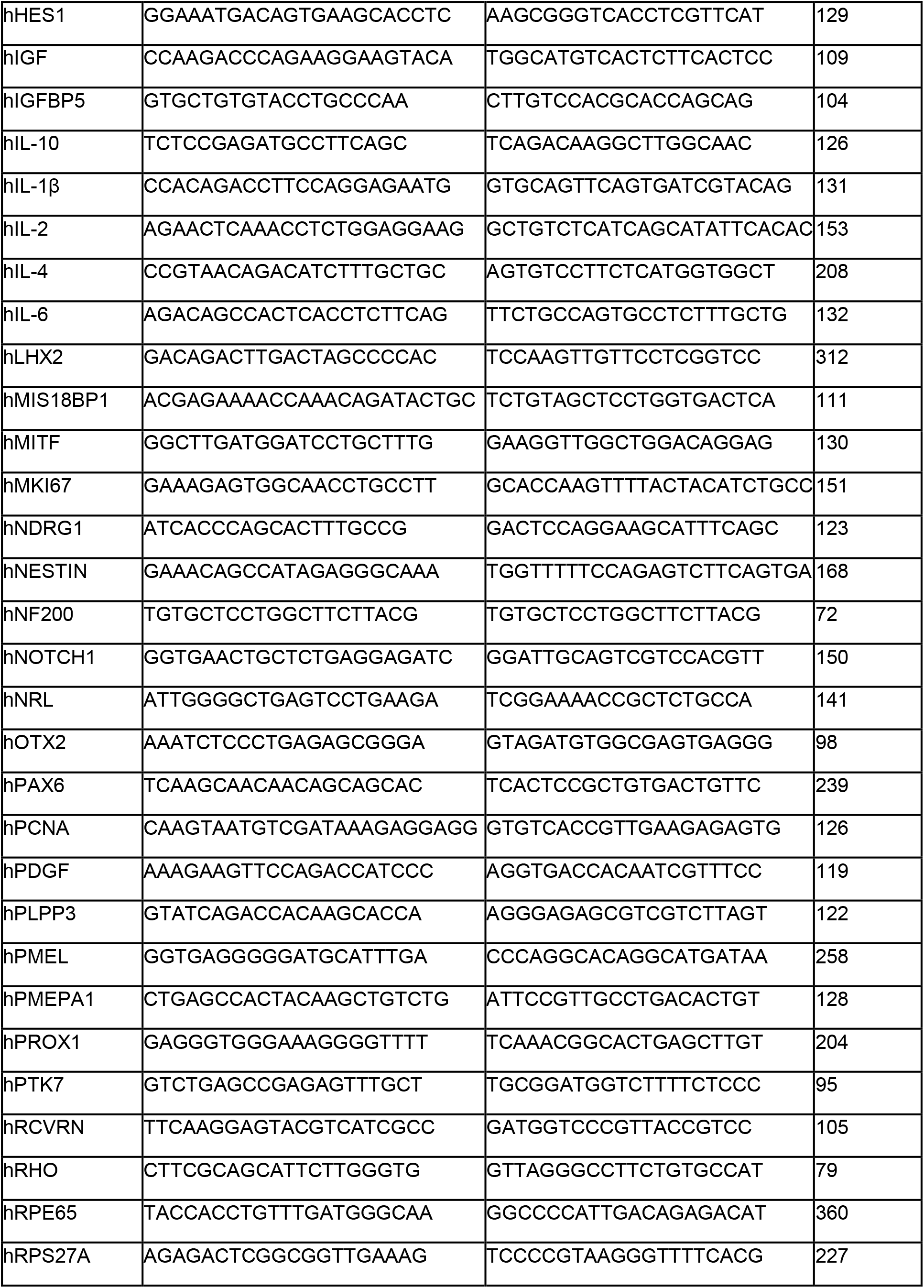
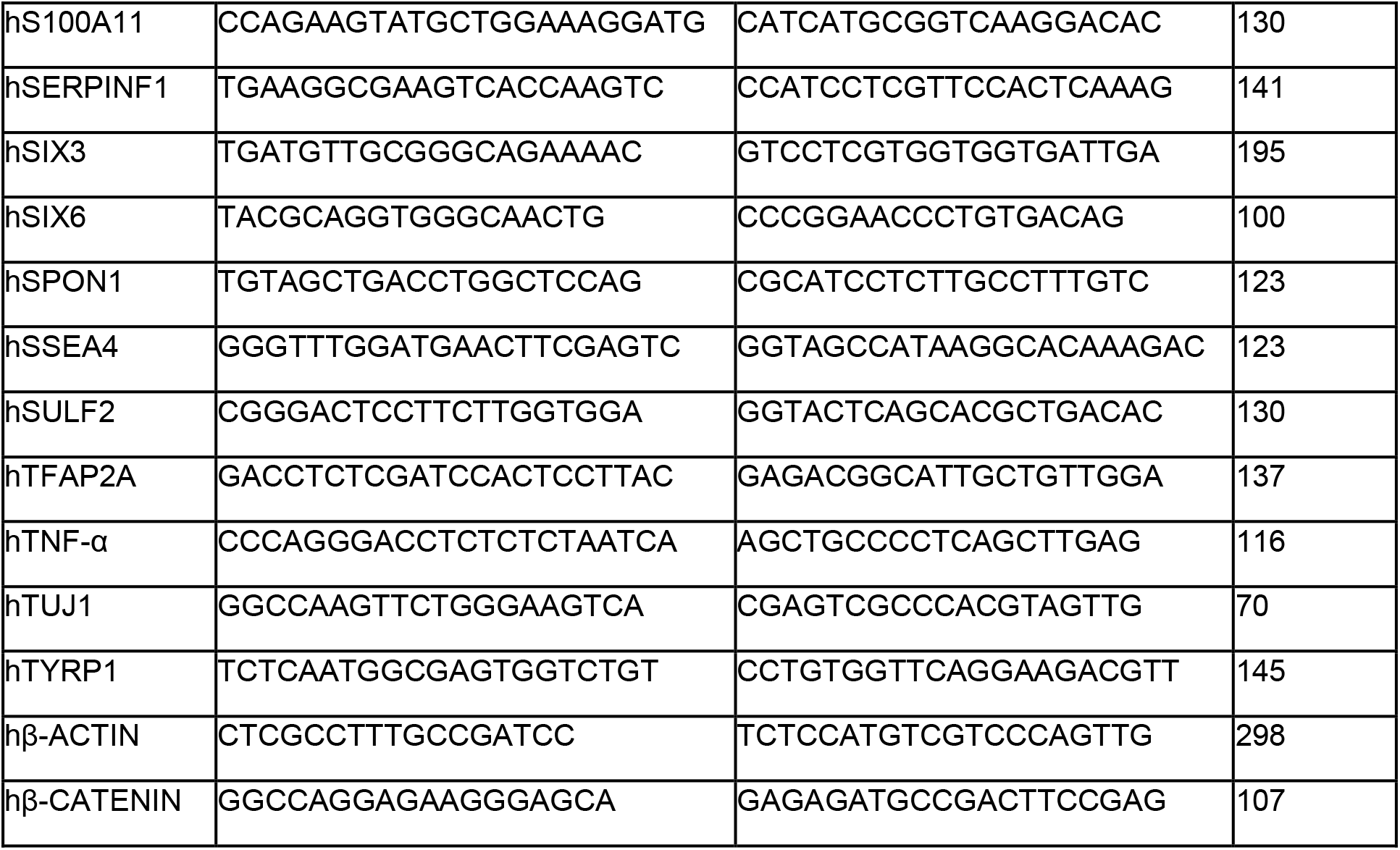
List of human primer sequences used in qRT-PCR.

**Supplementary Table 2.**
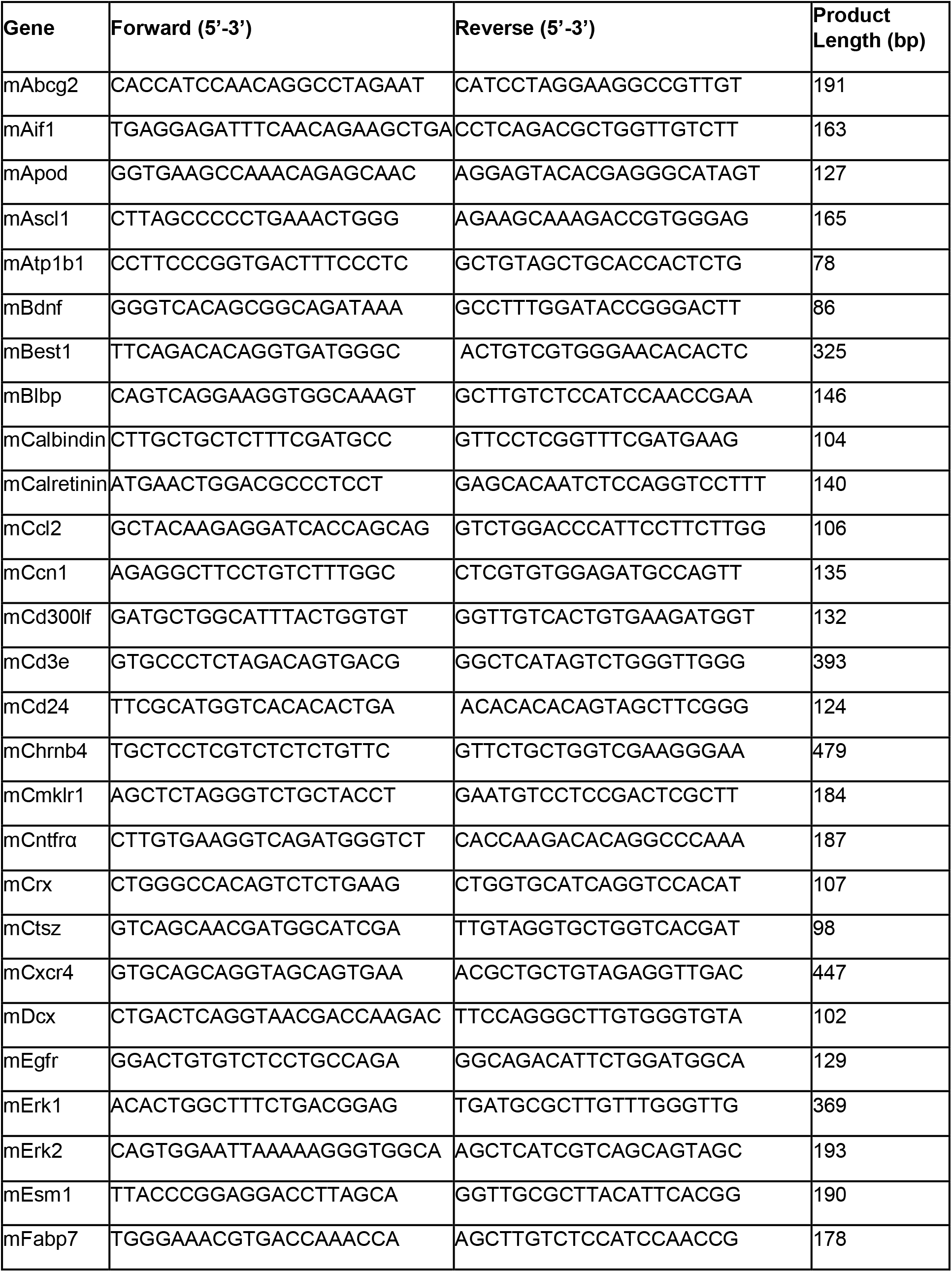
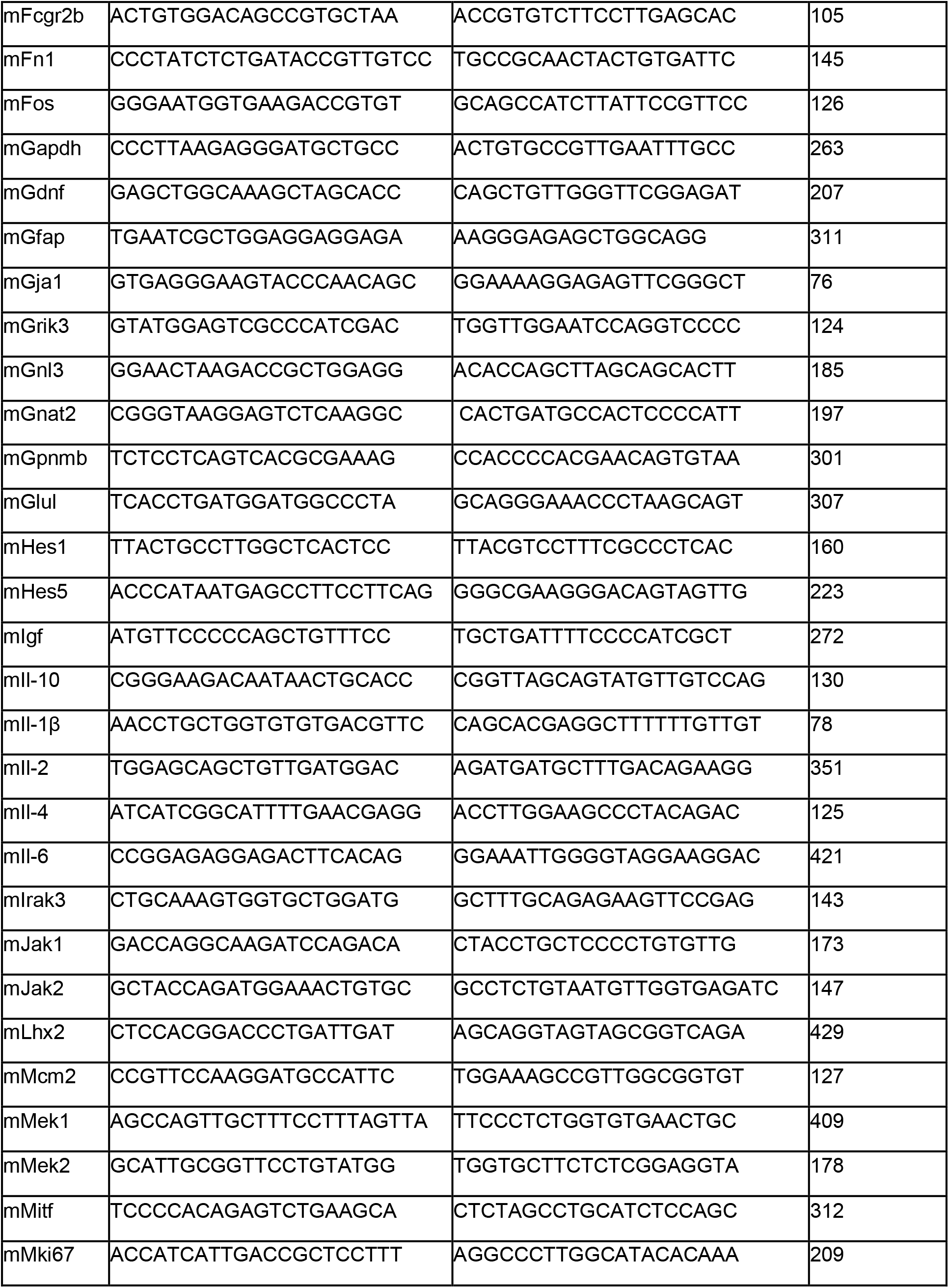
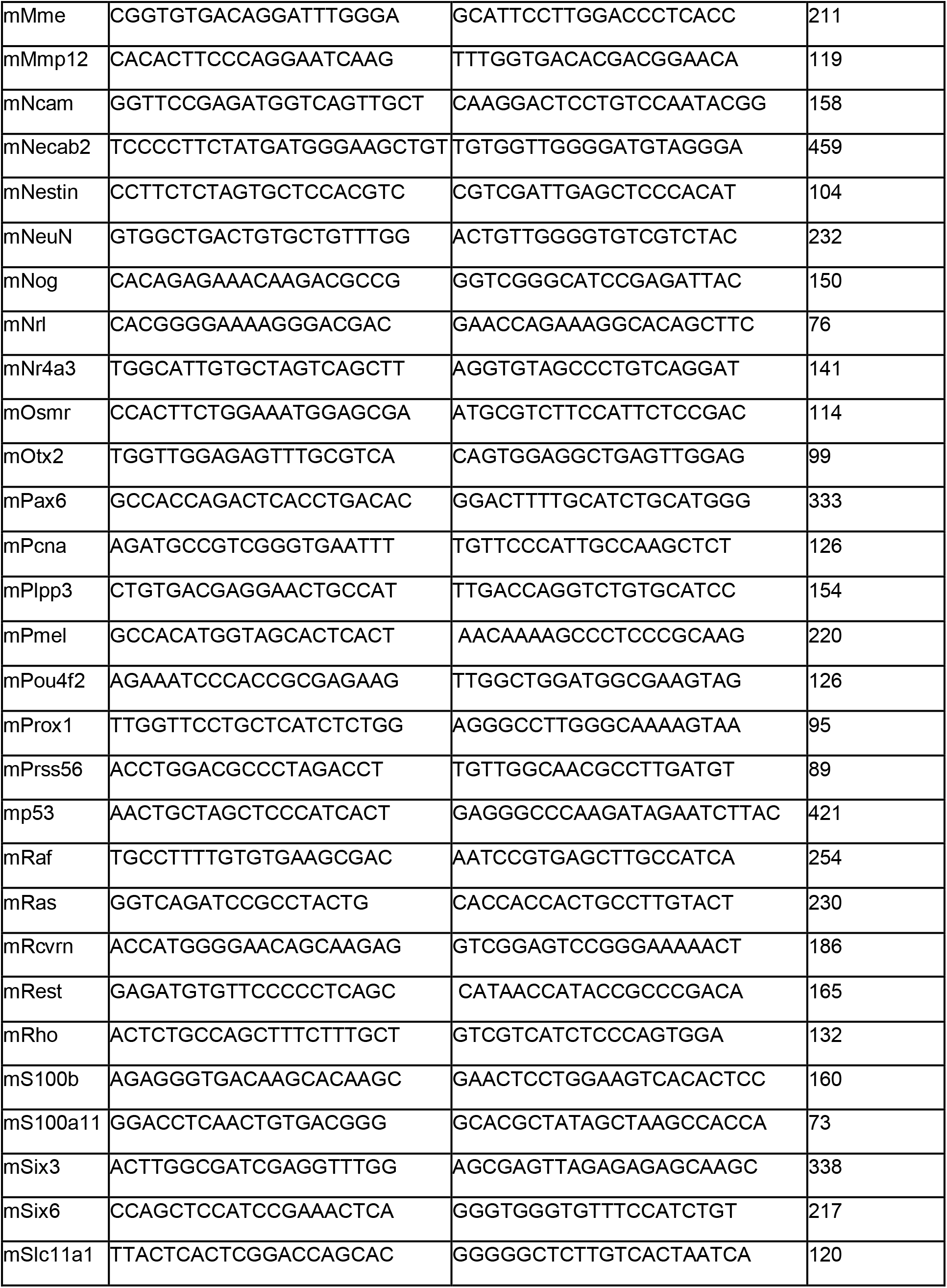
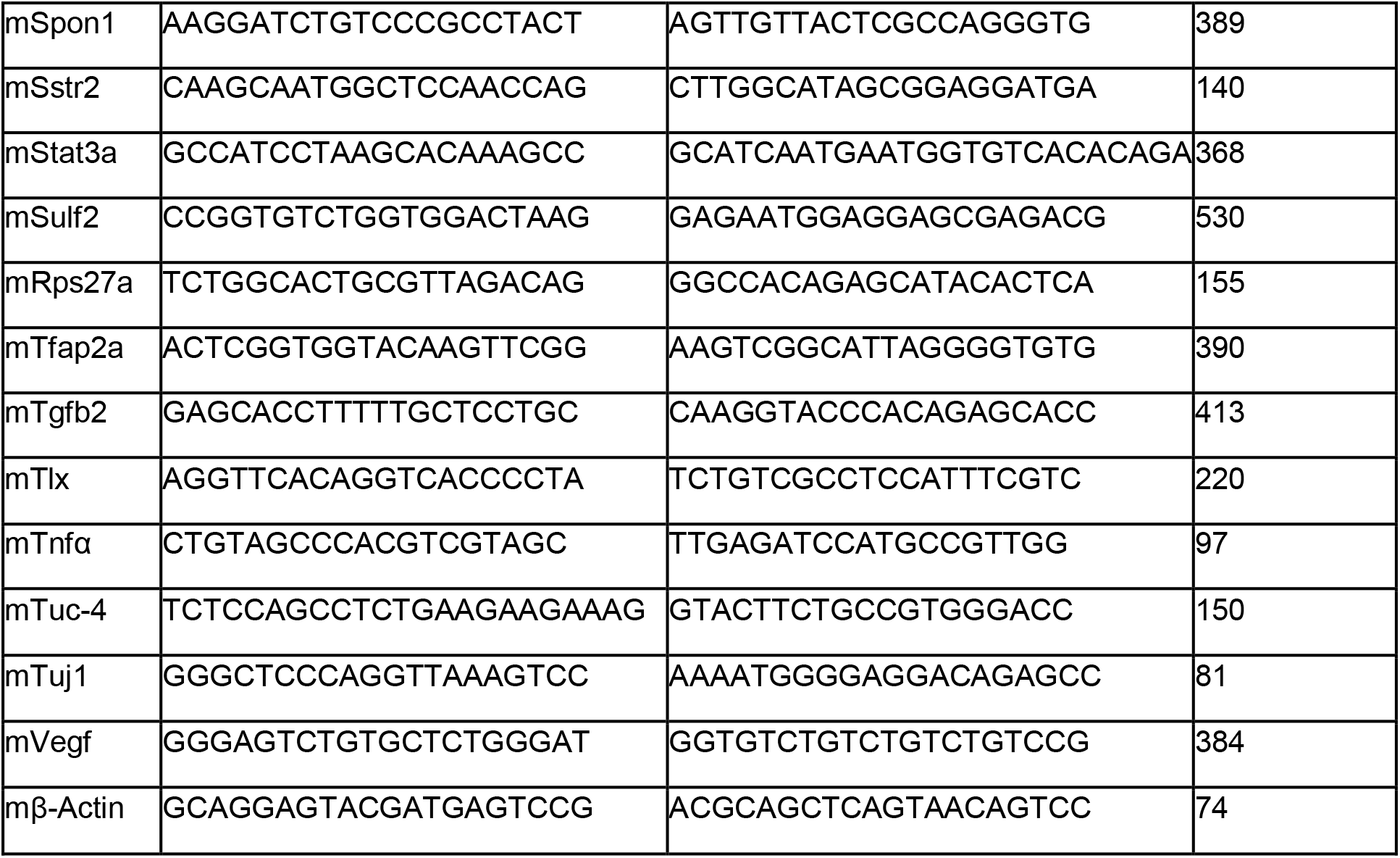
List of mouse primer sequences used in qRT-PCR.

**Supplementary Figure 1:**
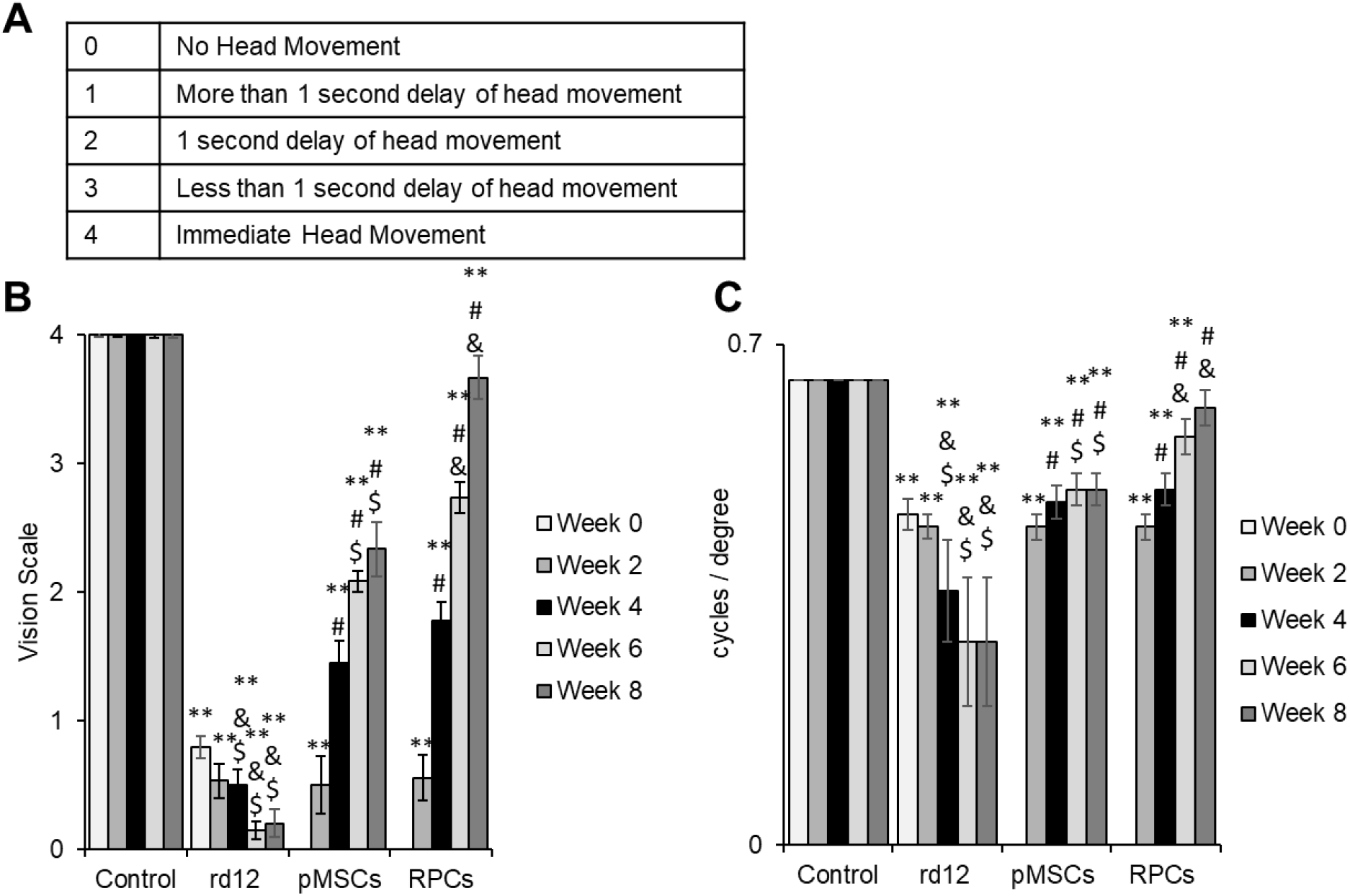
Behavioral analysis of mice. (A) Visual recovery scale. (B and C) Graphical representation of visual recovery and acuity assays at week 0, 2, 4, 6, 8 with WT, rd12, rd12+pMSCs and rd12+RPCs (**p ≤ 0.01). Symbols, **, #, & and $ indicate significant difference at p ≤ 0.01 between all experimental conditions: control (wild-type), rd12, rd12+pMSCs and rd12+RPCs, respectively.

**Supplementary Figure 2:**
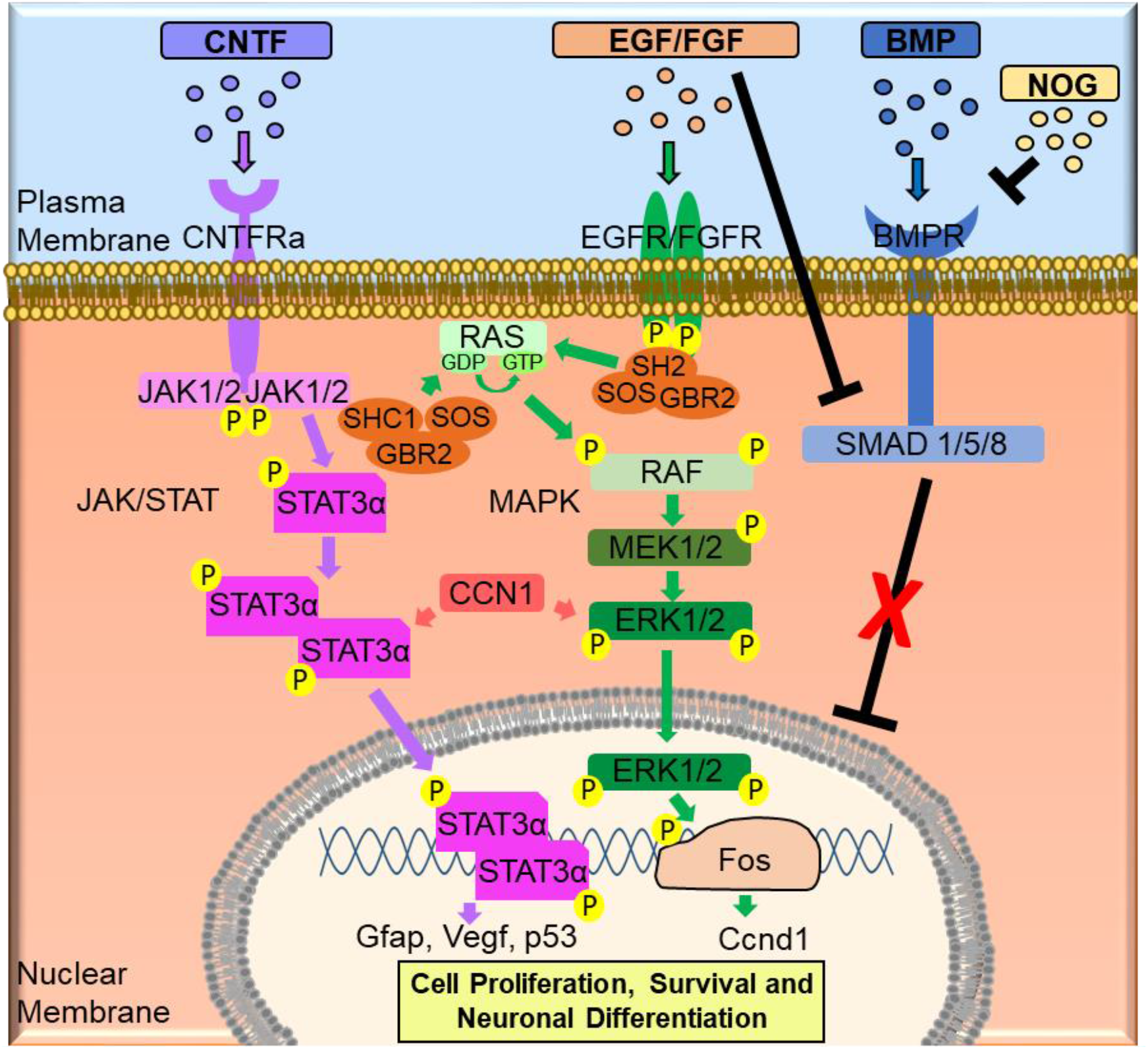
Proposed molecular mechanism involved the neuroprotection and retinal regeneration in rd12 mice transplanted with RPCs.

